# A conformational switch governs the dual role of liquid–liquid phase separation in amyloid fibrillization

**DOI:** 10.64898/2026.09.12.751119

**Authors:** Tianchen Li, Brayden Williams, Qi Han, Megan Steain, Margaret Sunde, Yi Shen

## Abstract

Biomolecular condensates provide dynamic environments that store and organize proteins, yet the fundamental principles determining whether condensation promotes or suppresses irreversible protein aggregation remain unclear. Here, we show that the receptor-interacting protein kinase 3 (RIPK3) RHIM domain follows distinct self-assembly pathways into condensates or amyloid fibrils depending on its conformational state. We find that predominantly folded proteins undergo liquid–liquid phase separation (LLPS), forming reversible condensates that kinetically suppress fibril formation. Partial unfolding instead promotes direct fibrillization that bypasses LLPS, whereas induced condensate formation via increasing ionic strength or addition of molecular crowder delays amyloid formation. In contrast, when the protein is predominantly unfolded, LLPS accelerates fibrillization through condensate-interface-mediated nucleation. These findings establish protein conformation as a determinant of whether condensates suppress or promote amyloid assembly, revealing a dual role for phase separation in regulating functional amyloid formation and providing a framework that connects condensate dynamics with cellular functions.

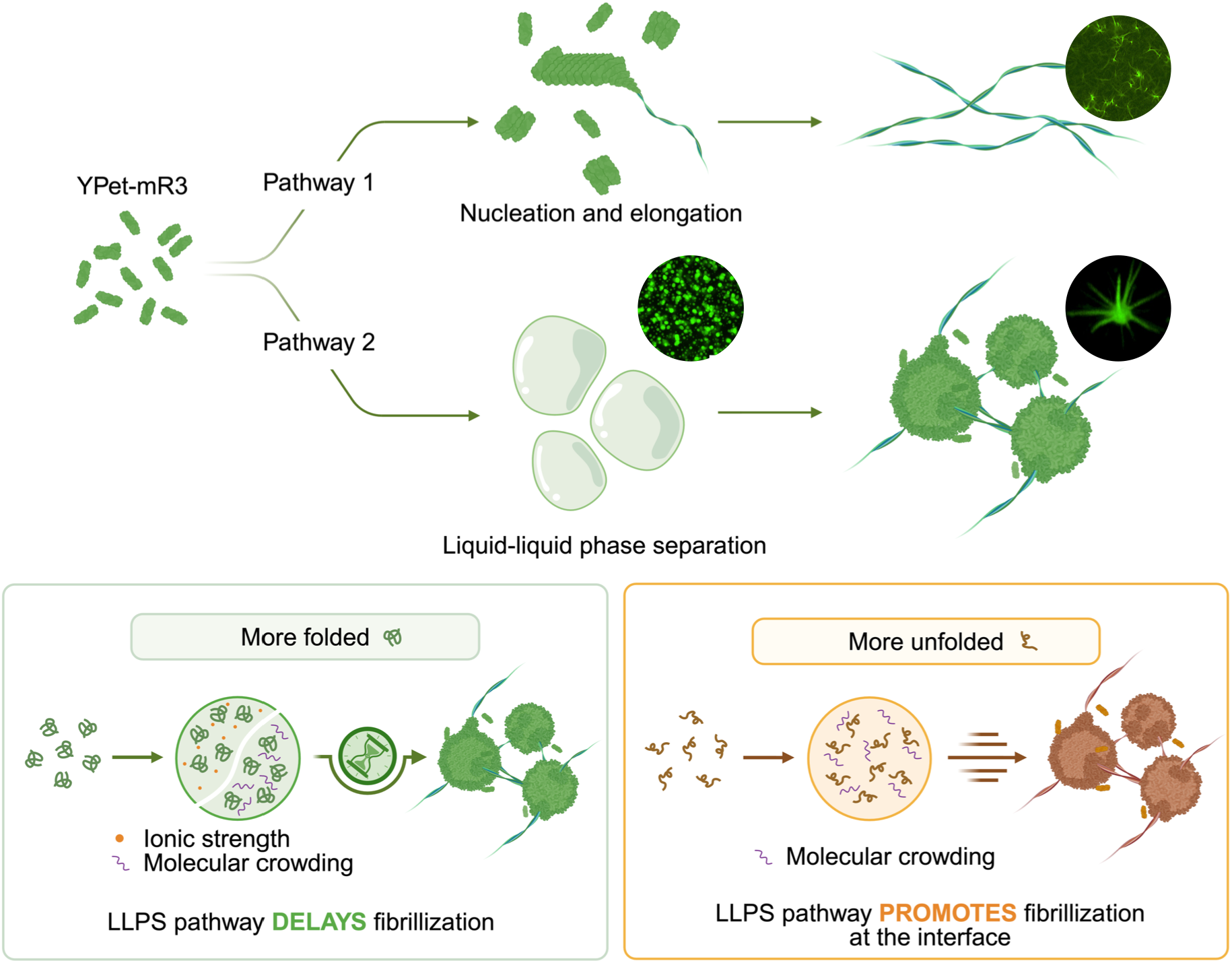

## Introduction

Necroptosis is an inflammatory programmed cell death pathway that is activated in response to infection or cellular damage. RIPK3 (Receptor-Interacting Protein Kinase 3) is a central protein downstream of all three of the necroptosis-initiating receptors, which include death receptors such as tumour necrosis factor (TNF) Receptor 1, Toll-Like Receptor 3 and 4 (TLR3/4), and Z-DNA Binding Protein 1 (ZBP1). RIPK3 functional amyloid fibril formation promotes self-phosphorylation and phosphorylation of its substrate MLKL. ^1^ Activated MLKL oligomerizes and is associated with rupture of the cell membrane, leading to inflammatory, lytic cell death. ^2,3^

Cells respond to stimulation of intrinsic sensors of danger by forming higher-order signalling complexes (signalosomes). In TNF-induced necroptosis, RIPK1 and RIPK3 co-assemble into a higher-order heteromeric functional amyloid signalling complex known as the classical necrosome. ^4^ The Z-form nucleic acid sensor ZBP1 binds Z-conformation nucleic acid and recruits RIPK3 to form a non-canonical necrosome. ^5^ Downstream of TLR3/4 activation, TRIF interacts with RIPK3 in a similar manner. ^6^ The recruitment and co-assembly of RIPK1, TRIF or ZBP1 with RIPK3, and the homotypic interactions of RIPK3, are driven by interactions between Receptor-interacting protein Homotypic Interaction Motifs (RHIMs) within these proteins. The RHIM is an ∼18-amino-acid sequence with high levels of conservation, particularly in the core tetrad (V/I-Q-V/I/L/C-G). ^7^ Previous studies have shown that the RHIM domain of murine RIPK3 readily self-assembles into amyloid fibrils, supporting the prevailing view that RHIM-mediated signalling is intrinsically amyloid-driven. ^7–9^

Biomolecular condensates have been studied for their essential roles in diverse cellular processes, including gene regulation, stress responses, and maintenance of cellular homeostasis. ^10^ Triggered by intrinsic factors, such as disease-associated mutations and post-translational modifications, or extrinsic cues, including changes in pH, ionic strength, and mechanical stress, condensates can undergo further liquid-to-solid transitions (LSTs), sometimes leading to irreversible protein aggregation. ^11^ Such transitions have been implicated in the pathogenesis of numerous neurodegenerative diseases, including FUS ^12^, hnRNPs ^13^, TDP-43 ^14^, α-synuclein ^15^, and tau ^16^. Conversely, other evidence suggests that condensates can also act as metastable reservoirs that delay amyloid formation by kinetically trapping proteins in a condensed but dynamic state. ^17^ Understanding the conditions under which condensate formation promotes or suppresses amyloid assembly therefore remains a fundamental unresolved question in protein phase transitions.

Emerging evidence indicates that LLPS also contributes to necroptotic signalling. OASL (2′-5′ oligoadenylate synthetase-like protein) can undergo LLPS and effectively recruit RIPK3 and ZBP1 to promote amyloid fibril formation and RIPK3 autophosphorylation. ^18^ Additionally, ZBP1 has been reported to form condensates during DNA- and RNA-virus infection. The Z-DNA binding domain of ZBP1 drives the LLPS, while the RHIM domains make the condensate more solid-like due to the formation of the amyloid-like structure. ^19^ Nevertheless, it remains unclear whether the RIPK3 RHIM domain itself can undergo LLPS and, more importantly, how LLPS influences RHIM amyloid nucleation and growth.

In this study, we demonstrate that YPet-tagged murine RIPK3 RHIM (YPet-mR3) (Fig. 1a) can self-assemble into amyloid fibrils through distinct pathways that either involve or bypass LLPS. YPet-mR3 undergoes LLPS at low concentrations of the denaturant urea, where it remains in a relatively compact conformation without progressing rapidly into irreversible fibrils. In contrast, at intermediate denaturant concentrations, partially unfolded YPet-mR3 bypasses LLPS and assembles directly into fibrils. The formation of irreversible fibrils could be delayed by increasing ionic strength or molecular crowding, which sequesters protein molecules into reversible condensates. However, at higher denaturant concentrations, when YPet-mR3 is even more unfolded, the LLPS pathway conversely accelerates fibril formation at the interface of condensates. These results demonstrate the dual role of condensates in protein self-assembly. Depending on protein conformational state and solution conditions, condensates can either act as reversible reservoirs that kinetically suppress amyloid formation or serve as catalytic interfaces that accelerate amyloid nucleation and fibril growth. Thus, in addition to roles in disease-associated amyloid formation, condensates may also play a role in the assembly of functional amyloids with consequences for cell signalling and cell death pathways.

**Figure 1.**
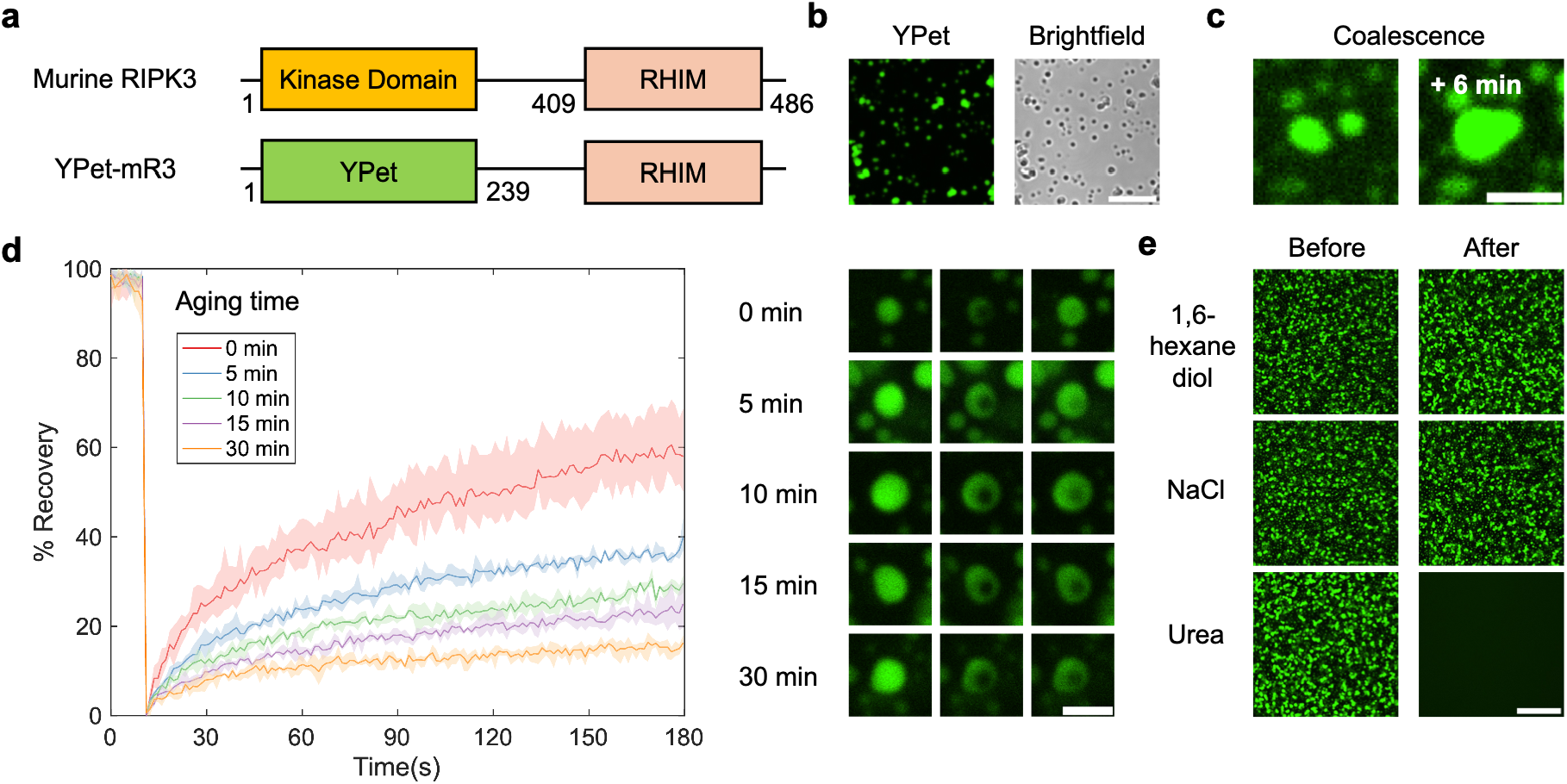
**a** Domain organisation of full-length murine RIPK3 protein and the YPet-mR3 construct used in this study. **b** Fluorescent and brightfield images of YPet-mR3 condensates. Scale bar: 10 μm. **c** Coalescence of YPet-mR3 condensates. Scale bar: 5 μm. **d** FRAP assay on YPet-mR3 condensates incubated at different time points. The error bars were measured based on three individual experiments. Scale bar: 2 μm. **e** The solubility of YPet-mR3 condensates incubated 1 hour after mixing with 20 wt% 1,6-hexanediol, 1 M NaCl, and 8 M Urea. Scale bar: 20 μm.

## Results

### LLPS of YPet-mR3 *in vitro*

Previous studies showed that YPet-mR3 gradually assembles into amyloid fibrils when the denaturing buffer (such as guanidium chloride or urea) was removed by dialysis. ^7,8^ In our experiments, rather than using dialysis, YPet-mR3 was diluted 10-fold from the stock solution containing 8 M urea. To our surprise, YPet-mR3 formed spherical, liquid-like droplets within 2 minutes (Fig. 1b), in contrast to the extended fibrillar structure previously reported under dialysis conditions. ^8,20^ The neighbouring newly formed droplets can contact and coalesce, although fusion was frequently incomplete, suggesting restricted fluidity or rapid structural maturation (Fig. 1c). The dynamics of protein molecules inside the condensates were assessed using fluorescence recovery after photobleaching (FRAP) (Fig. 1d). Freshly formed condensates recovered around 60% in the photobleached area within 180 seconds. A rapid reduction in dynamicity was observed, with the recovery rate decreasing from 60% to 16% within 30 minutes. This time-dependent dynamical arrest resembles the aging behavior reported for yeast prion protein condensates, whose FRAP recovery decreases from 100% to less than 60% in one hour. ^21^

Biomolecular condensates are stabilized by multiple types of molecular interactions, such as hydrogen bonding, ionic interactions, hydrophobic interactions, aromatic interactions, etc. ^11^ To probe the driving force of LLPS, we destabilized the condensates with reagents that perturb different classes of intermolecular interactions (Fig. 1e). The condensates can be fully dissolved by mixing with 8M urea, indicating LLPS depends on interactions or conformational states which are sensitive to partial denaturation. A similar urea sensitivity has been reported for GK-16 peptides inspired by the mussel foot protein, in which LLPS is mediated by the hydrogen bonding between Dopa (L−3,4-dihydroxyphenylalanine, a post-translationally modified form of tyrosine) residues. ^22^ By contrast, YPet-mR3 condensates cannot be dissolved by mixing with 1,6-hexanediol (20 wt%) or NaCl (1 M), suggesting that hydrophobic or electrostatic interactions may not be the dominant driving forces behind LLPS. ^23,24^

### Urea and salt mediate YPet-mR3 phase transitions

We noticed that changing the urea concentration allowed us to modulate the phase separation of YPet-mR3. Thus, to map the phase diagram of YPet-mR3 LLPS, the effect of YPet-mR3 protein concentration was investigated across a gradient of urea (Fig. 2a and Supplementary Fig. S1a). At 0.8 M urea, YPet-mR3 can undergo LLPS without the addition of salt. At 1.4 M urea, when protein concentration was below or equal to 10 μM, YPet-mR3 formed irregular fibrils with less defined morphology after 24-hour incubation, without LLPS being observed (Supplementary Fig. S1a). When YPet-mR3 was above 10 μM, YPet-mR3 favoured condensate formation rather than fibrillization. At 2 M urea, YPet-mR3 formed clearly defined fibrils without LLPS across all concentrations examined. No phase transitions occurred within 24 hours when the urea concentration exceeded 3 M, or the protein concentration was below 1 μM. (Supplementary Fig. S1a) These observations show that the selectivity between LLPS and direct fibrillization depends on both protein concentration and the conformational perturbation induced by urea.

**Figure 2.**
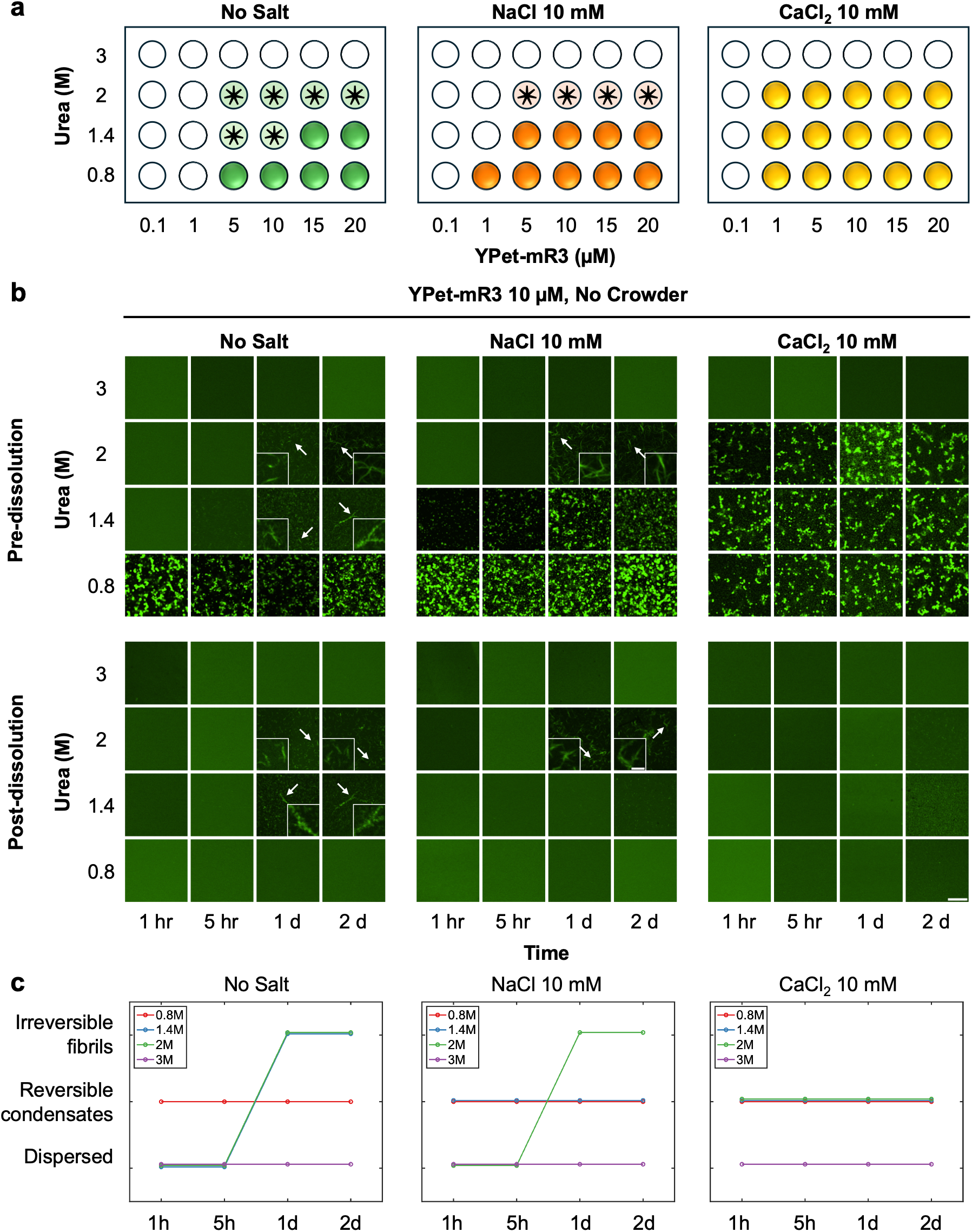
**a** Phase diagram of YPet-mR3. The highlighted regions indicate that LLPS was observed. The areas labelled with asterisks indicate that fibrillization without LLPS was observed. **b** Confocal microscopic images of YPet-mR3 incubated at different time periods and after dissolution. Scale bar: 10 μm. Insert: 2 μm. Three independent replicates were performed, showing similar results. **c** The reversibility of YPet-mR3 phase transitions from (b). The addition of salts effectively delayed the irreversible fibrillization under 1.4 M and 2 M urea.

Ionic strength can modulate protein phase transitions by screening the charge on protein molecules. We added NaCl and CaCl_2_ to assess if monovalent and divalent ions can affect YPet-mR3 phase transitions. The addition of NaCl and CaCl_2_ was found to promote LLPS of YPet-mR3. Both salts expanded the range of conditions supporting YPet-mR3 LLPS, with CaCl_2_ producing a more pronounced shift in the LLPS boundary. (Fig. 2a and Supplementary Fig. S1b, c). The addition of salt also altered the competition between LLPS and fibrillization. At 1.4 M urea, YPet-mR3 underwent LLPS in the presence of NaCl or CaCl_2,_ and no subsequent fibrillization was observed within 24 hours. At 2 M urea, NaCl did not induce detectable LLPS, and YPet-mR3 formed fibrils after 24 hours. By contrast, CaCl_2_ redirected YPet-mR3 towards condensate formation. MgCl_2_ similarly promoted LLPS at 2 M urea (Supplementary Fig. S2), suggesting that divalent binding facilitates the LLPS of YPet-mR3. This behavior is consistent with previous observations that divalent cations can promote the LLPS of α-synuclein. ^25^

To further investigate how condensates can modulate irreversible aggregate formation, we prepared YPet-mR3 condensates at 10 μM, a concentration comparable to that used in previously reported studies ^7,18^. The samples were incubated under different urea and salt concentrations for defined time periods, then mixed with 8 M urea (Fig. 2b,c). The reversible condensates were expected to dissolve under these denaturing conditions, whereas clearly defined or irregular fibrillar assemblies were expected to remain urea-resistant. ^4^ At 0.8 M urea, condensates formed within 48 hours under all salt conditions remained fully soluble. The fibrils formed at 1.4 M and 2 M urea without salt, and at 2 M urea with NaCl after 24 hours of incubation resisted dissolution in urea. By contrast, the condensates incubated for 48 hours in 1.4 M urea with NaCl and 1.4-2 M urea with CaCl_2_ could be fully dissolved.

To determine whether these condensates remained reversible over longer periods, samples were further incubated for up to 7 days (Supplementary Fig. S3). At 1.4 M urea with NaCl and 2 M urea with CaCl_2_, the condensates subsequently developed clearly defined or irregular fibrillar structures that were not fully dissolved by urea. In contrast, no fibrils were observed with YPet-mR3 condensates incubated at 0.8 M urea under any of the salt conditions examined, or at 1.4 M urea with CaCl_2_. The condensates could be dissolved by mixing with 8 M urea. These results show that YPet-mR3 condensates can remain urea-soluble despite the pronounced dynamical arrest, indicating that their intermolecular interaction networks remain reversible. Increasing ionic strength, particularly through the addition of divalent cations, can delay the irreversible fibril formation and temporarily trap YPet-mR3 as reversible condensates.

### Conformational changes of YPet-mR3 at different urea concentrations

Next, we investigated the mechanism of how urea concentration influences the conformational state of YPet-mR3 and thereby correlates with its phase behavior. Transmission electron microscopy (TEM) highlighted the distinct morphology between condensates formed with urea at 0.8 M and the fibrillar structures formed by YPet-mR3 in 2 M urea (Fig. 3a). Since urea is a commonly used denaturant that alters protein structure and folding, we examined its effect on YPet-mR3 conformation using intrinsic fluorescence spectroscopy. Increasing urea concentrations induced a progressive red shift in the intrinsic fluorescence emission maximum (Fig. 3b). This shift indicates increased solvent exposure of tryptophan residues and progressive disruption of the native protein conformation, consistent with previous studies of urea-induced protein unfolding. ^26,27^ Circular dichroism spectra also revealed a similar change in conformational states at increasing urea concentrations, indicated by the descending slope between 230 and 240 nm (Supplementary Fig. S4). ^28^ Time-resolved FTIR spectroscopy further indicated different secondary structure transitions within the first 5 hours. YPet-mR3 incubated under 0.8 M urea displays an Amide I peak near 1639 cm^-1^ and shifted to 1631 cm^-1^. By contrast, at 2 M urea, the initial Amide I peak was centred near 1647 cm^-1^, and subsequently shifted to 1633 cm^-1^, accompanied by the emergence of a shoulder peak near 1670 cm^-1^ (Fig. 3c). ^29^ These spectral changes are consistent with the progressive formation of β-sheet-rich structures under both conditions but with distinct pathways. At 0.8 M urea, LLPS of YPet-mR3 features a smoother transition into β-sheet-rich structure. By contrast, YPet-mR3 under 2 M urea displayed a less ordered conformation initially, followed by a more abrupt transition to a β-sheet-rich structure over time, consistent with the phenomenon of direct fibrillization (Supplementary Fig. S5).

**Figure 3.**
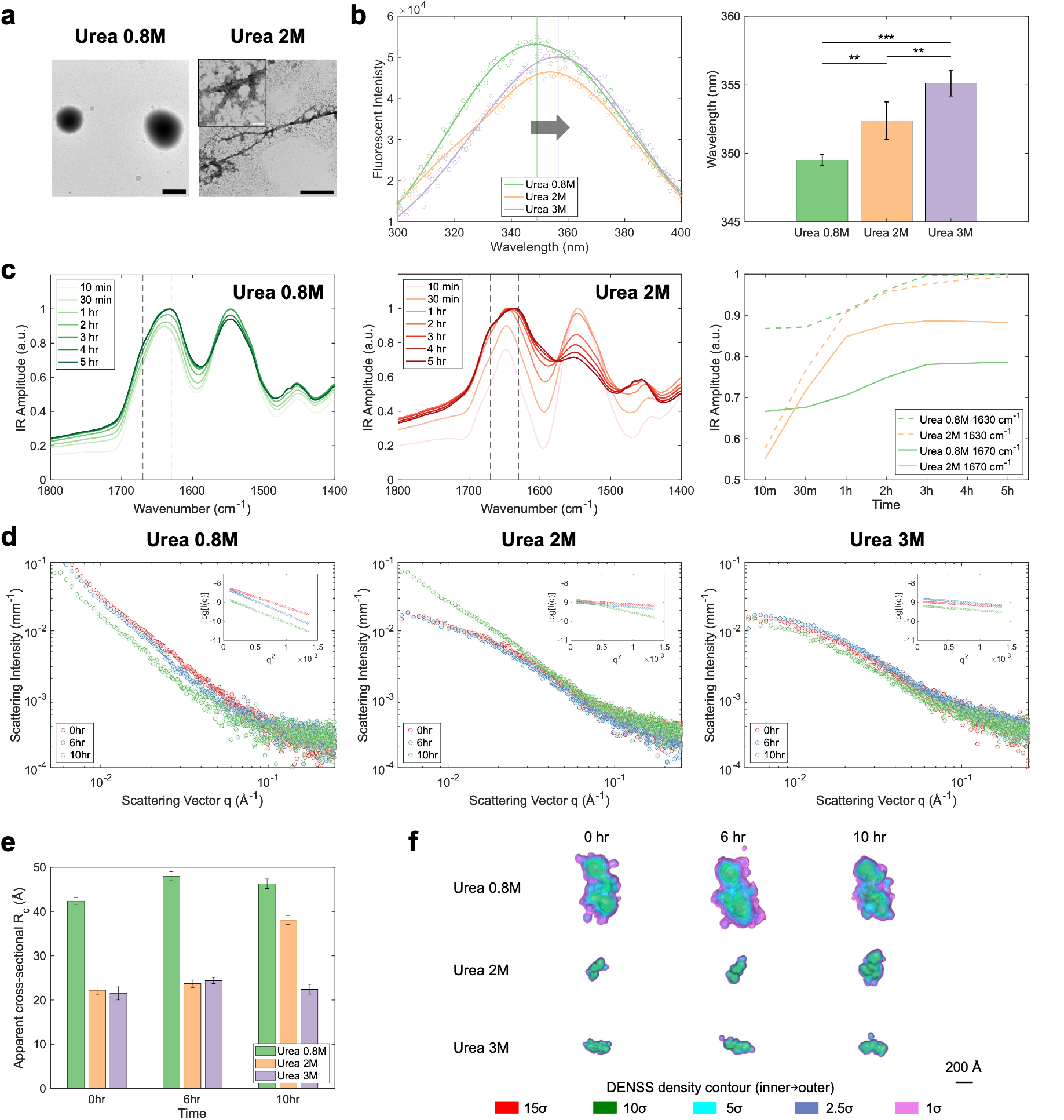
**a** TEM images of YPet-mR3 condensates formed under 0.8 M urea and fibrils formed under 2 M urea, in the presence of Tris-HCl 50 mM. Scale bar: 1 μm. Insert: 100 nm. **b** Intrinsic fluorescence spectra of YPet-mR3 in the presence of 0.8 M, 2 M, and 3 M urea. The bar plot represents the wavelengths of maximum fluorescence intensity from 4 biological replicates, with error bars indicating standard deviations. ** p < 0.01, *** p < 0.001. **c** FTIR spectra measured during YPet-mR3 LLPS under 0.8 M urea and fibrillization under 2 M urea. The time-dependent amplitude at 1630 cm^-1^ and 1670 cm^-1^ is shown on the right. **d** X-ray scattering intensity *I*(*q*) as a function of scattering vector *q*. Measurements were taken from 10 μM YPet-mR3 incubated under 0.8 M, 2 M and 3 M urea at 0, 6, and 10 hours. Insets: cross-sectional Guinier fits, ln[*q*·*I*(*q*)] versus *q*² over the fixed window *q* = 0.012–0.035 Å⁻¹. **e** The apparent cross-sectional radius of gyration *R_c_* from cross-sectional Guinier analysis (fixed window). **f** DENSS averaged electron-density envelopes of YPet-mR3 under 0.8 M, 2 M and 3 M urea at 0, 6, and 10 hours, all shown at a common scale (1700 Å field) and aligned to each condition’s 0 h reconstruction (four nested contour levels: 1.5σ, 2.5σ, 5σ and 10σ). Each map is the average of ten independent reconstructions. The 0.8 M envelopes are not uniquely determined (map-to-map Fourier shell correlation resolution comparable to the particle R_g_) and are shown for illustration only (Supplementary Fig. S10 and Table S1). Scale bar: 200 Å.

Small-angle X-ray scattering (SAXS) was performed on YPet-mR3 incubated under 0.8 M, 2 M and 3 M urea at 0, 6, and 10 hours (Fig. 3d). Because YPet-mR3 self-associates into large, polydisperse assemblies, the profiles do not display a classical Guinier region at any accessible *q*. All size parameters are therefore treated as apparent, intensity-weighted averages and used only for relative comparison. At 0.8 M urea, the intensity at low *q* is highest at all time points and the log–log *I*(*q*) profiles retain a similar shape over time, consistent with large, compact clusters that do not substantially elongate. At 2 M urea, the overall intensity is lower, and the low-*q* region is progressively enhanced with a subtle change in slope between the 6^th^ and 10^th^ hours, consistent with growth of the assemblies. To compare the effective thickness of these elongated and clustered assemblies while minimising the influence of length polydispersity and weak low-*q* aggregation, we applied a cross-sectional Guinier analysis, fitting ln[*q*·*I*(*q*)] versus *q*² over a single fixed window (*q* = 0.012–0.035 Å⁻¹) common to all samples, and extracted an apparent cross-sectional radius of gyration, *R_c_* (Fig. 3d insets and Fig. 3e). *R_c_* was ∼50 Å at 0.8 M urea (at the edge of cross-sectional-Guinier validity, *q*·*R_c_* ≈ 1.6, and therefore an effective value only), ∼20 Å at 2 M urea at the 0^th^ and 6^th^ hour, increasing to 30–40 Å at the 10^th^ hour, and ∼20 Å at 3 M urea at all time points; the relative ordering (0.8 M > 2 M ≈ 3 M) is robust to the choice of window (Supplementary Fig. S6). Dimensionless Kratky plots (Supplementary Fig. S7), used only qualitatively, show an upshift in the low-*qR_g_* region at the 10^th^ hour for 2 M urea, consistent with the appearance of short fibrils between the 6^th^ and 10^th^ hours (Supplementary Fig. S5). At 3 M urea, the curves nearly overlap over time with only a modest low-*q* excess, consistent with YPet-mR3 remaining dispersed.

The pair-distance distributions p(r) show the same trend in real space: broad and essentially unchanged over time at 0.8 M urea, broadening markedly between the 6^th^ and 10^th^ hour at 2 M urea, and narrow throughout at 3 M urea (Supplementary Fig. S8). Because the assemblies are polydisperse and, at 0.8 M urea, larger than the size range strictly resolved by the accessible q, p(r) and the derived D_max_ are used as relative indicators of size only (Supplementary Table S1). These results indicate that, under 0.8 M urea, relatively folded YPet-mR3 forms clusters resistant to elongation into an amyloid-like structure, associated with LLPS, ^30^ whereas fibril formation is favoured when YPet-mR3 is slightly more unfolded (2 M) and is suppressed when it is even more unfolded and remains dispersed (3 M). Accordingly, ab initio shape reconstructions, based on the p(r), at the 0^th^ hour were computed both as bead models (DAMMIF; Supplementary Fig. S9) and as electron-density maps (DENSS; Fig. 3f and Supplementary Fig. S10). At 0.8 M urea the reconstruction is a large envelope consistent with higher-order clustered assemblies; at 2 M urea it is smaller and more elongated; and at 3 M urea smaller still, in agreement with the *R_c_* and p(*r*) analyses. These envelopes are low-resolution and, for the 0.8 M clusters, not uniquely determined (independent reconstructions did not converge; DENSS map-to-map Fourier shell correlation ≈ 230 Å, comparable to the particle *R_g_*); they are therefore interpreted qualitatively (Supplementary Figs S6–S11 and Tables S1 and S2).

### The role of core tetrad residues on YPet-mR3 phase transitions

The core tetrad residues VQIG contribute to a key β-strand within murine RIPK3 amyloid fibrils, and mutation of VQIG disrupts TNF-induced necroptosis and suppresses cell death. ^8,31^ We therefore assessed how VQIG contributes to YPet-mR3 LLPS and fibrillization by mutating the VQIG residues in YPet-mR3 to AAAA (YPet-mR3-mut) (Fig. 4a). YPet-mR3-mut is still capable of LLPS under 0.8 M urea, while it is arrested more quickly compared to the wild type (Fig. 4b). The fresh YPet-mR3-mut condensates display a 40% recovery after photobleaching, whereas the fresh wild-type condensates can reach 60% recovery. The recovery rate decreases to 15% after 10 minutes and to 6% after 30 minutes, whereas the wild type drops only to 30% and 16%, respectively. YPet-mR3-mut condensates can be dissolved by urea but not by NaCl or 1,6-hexanediol, implying that the driving force of LLPS does not differ from that of the wild type (Fig. 4c). YPet-mR3-mut can form condensates across 0.8M – 2M urea and the AAAA mutation results in loss of the capability of forming fibrils (Fig. 4d, e). None of the conditions generated a fibrillar structure even after 7 days of incubation. Instead, they form a pearl-chain-like structure consisting of aged condensates (Supplementary Fig. S12). These results highlight that while the core tetrad residues are crucial for amyloid formation, they may not be the critical multivalent binding sites for condensate formation.

**Figure 4.**
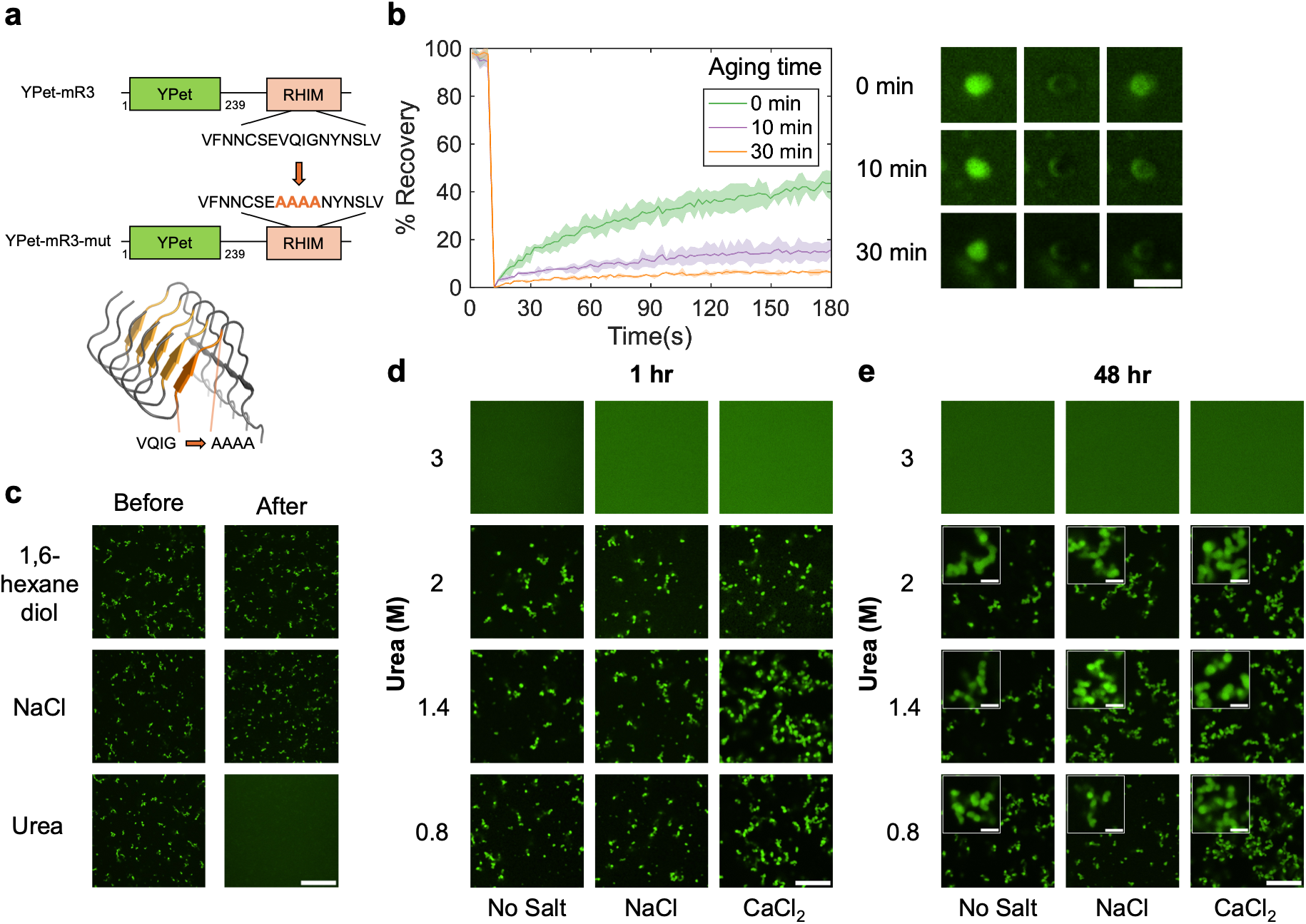
**a** The construct of YPet-mR3-mut. The core-tetrad residues that contributed to the amyloid core (PDB: 6JPD) are mutated. **b** FRAP assay on YPet-mR3-mut condensates incubated at different time points. The error bars were measured based on three individual experiments. **c** The solubility of YPet-mR3-mut condensates incubated 1 hour after mixing with 20 wt% 1,6-hexanediol, 1 M NaCl, and 8 M Urea. Scale bar: 20 μm. **d,e** Confocal microscopic images of YPet-mR3-mut incubated under different conditions after 1 hour (**d**) and 48 hours (**e**). Scale bar: 10 μm. Inserts: 2 μm.

### Modulating the phase behavior of YPet-mR3 by using molecular crowder

To mimic a crowded intracellular environment, molecular crowders such as PEG, Dextran, and Ficoll are commonly used for *in vitro* studies. It has been found that molecular crowding can usually enhance LLPS. This can be due to the increase in the local concentration of protein molecules, changes in the protein conformation and the degree of hydration. ^32^ We therefore aimed to determine whether adding a molecular crowder could enhance LLPS and alter the phase behavior of YPet-mR3.

Addition of PEG8000 markedly expanded the range of conditions under which YPet-mR3 underwent LLPS. In the presence of 10 wt% PEG8000, YPet-mR3 formed condensates from 0.8 M to 5 M urea (Fig. 5a, b, and Supplementary Fig. S13), substantially extending the LLPS regime observed in the absence of crowder. In the presence of both PEG8000 and CaCl_2_, YPet-mR3 forms LLPS even at 6 M urea (Supplementary Fig. S13). PEG8000 also significantly lowers the critical phase-separation concentrations. The critical separation concentration (C_sat_) of YPet-mR3 condensates was measured in the presence of 10 wt% PEG8000 via centrifugation (Fig. 5c). ^33^ The presence of molecular crowder had little impact on C_sat_ under 0.8 M urea, which was 1.27 μM without crowder and 1.20 μM with PEG8000. By contrast, in the presence of molecular crowder, the C_sat_ are measured as 1.64 μM and 2.67 μM at 2 M and 3 M urea, respectively, whereas under these conditions YPet-mR3 does not phase separate at 10 μM in the absence of crowder. It was reported that crowding reagents can either remain in the diluted phase ^12,34^ or partition into the condensate phase. ^35,36^ To verify how PEG is distributed in the system, we used a mixture of 1 wt% Cy5-tagged PEG8000 and 9 wt% non-tagged PEG8000. It was found that Cy5-tagged PEG8000 was strongly recruited into condensates across all urea concentrations tested (Supplementary Fig. S14). The co-condensation of PEG8000 and YPet-mR3 increases the local concentration of protein molecules, thereby enabling phase separation under high-urea conditions. ^35^

**Figure 5.**
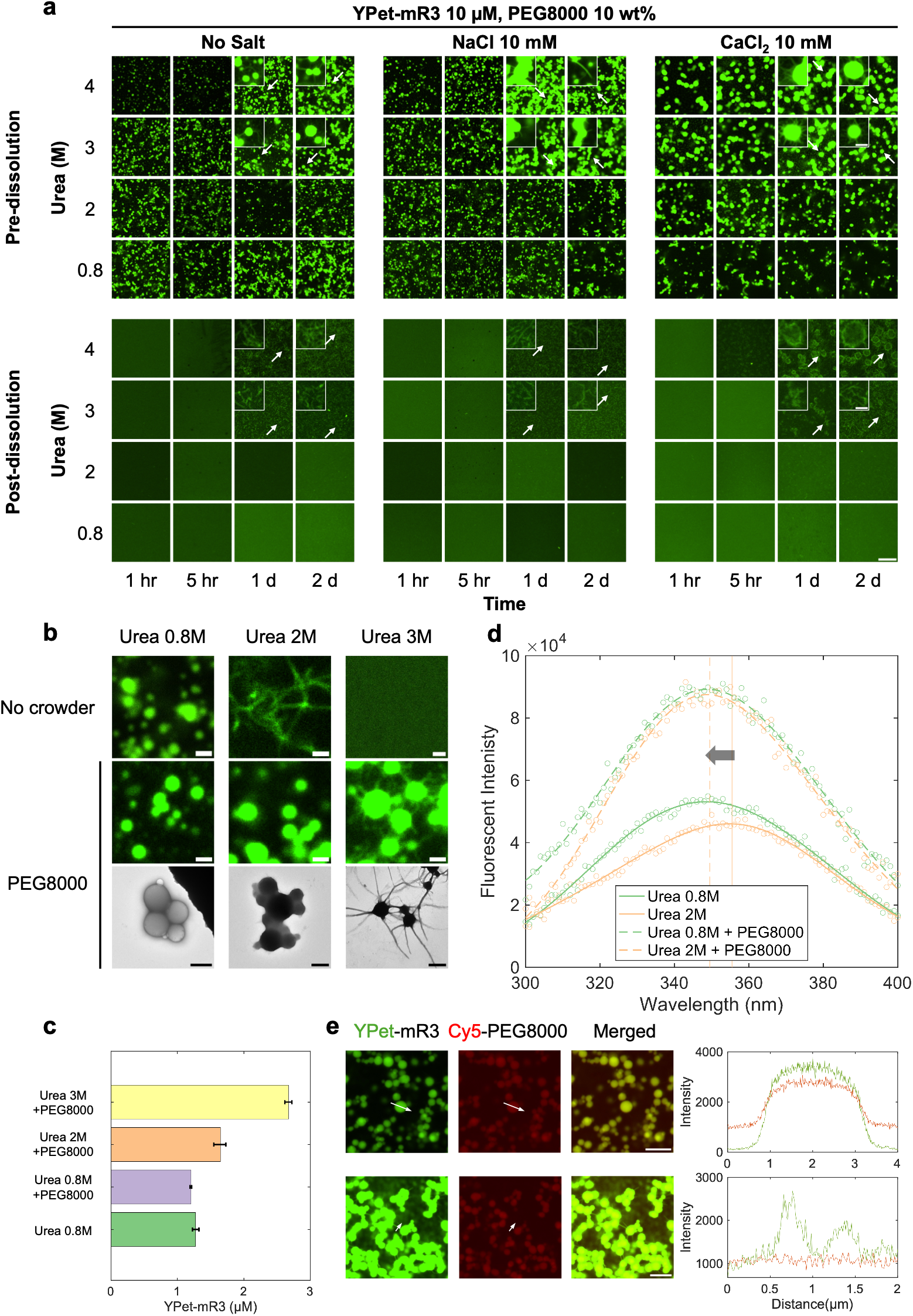
**a** Confocal microscopic images of YPet-mR3 incubated at different time periods in the presence of 10 wt% PEG8000 and after dissolution. Scale bar: 10 μm. Zoom-in images: 2 μm. Three independent replicates were performed, showing similar results. **b** Top: Confocal images of 10 μM YPet-mR3 incubated at different urea concentrations without molecular crowding. Bottom: Confocal microscopic and TEM images of 10 μM YPet-mR3 incubated at different urea concentrations with 10 wt% PEG8000. Scale bar: 1 μm. **c** Critical separation concentration of YPet-mR3 under different urea concentrations in the absence or presence of 10 wt% PEG8000. The error bar represents 3 biological replicates. **d** Intrinsic fluorescence spectra of YPet-mR3 under 0.8 M and 2 M urea with/without 10 wt% PEG8000. **e** Confocal microscopic images of YPet-mR3 incubated 24 hours under 3 M urea in the presence of 10 wt% PEG8000 (1wt% Cy5-PEG8000 + 9wt% PEG8000) and 50 mM Tris-HCl. In the second row, the YPet channel is overexposed to visualize the fibril. The intensity profile of the labelled condensate and fibrils is shown. Scale bar: 10 μm.

The role of crowder-induced LLPS in fibrillization depended strongly on urea concentration and, by inference, on the conformational states of the protein. At 2 M urea, the addition of PEG8000 suppressed fibril formation over 48 hours, whereas irreversible fibrils were formed under no-salt conditions or with NaCl (Fig. 5a, b). Under these conditions, molecular crowding redirected YPet-mR3 into condensates and delayed fibril formation. As we have previously shown that the folding states influence the distinct phase transition pathways, we measured the intrinsic fluorescence of YPet-mR3 in the presence of PEG8000 (Fig. 5d). The position of maximum fluorescence intensity at 0.8 M urea remained unchanged. At 2 M urea, the maximum fluorescence intensity shifted from 355.5 nm to 349.5 nm. This blue shift indicates the reduced solvent exposure of the tryptophan residues and a more folded conformation.

However, at 3 M and 4 M urea, fibrils were observed growing on the interface of the condensates, producing characteristic sea-urchin-like morphologies (Fig. 5a, b, and Supplementary Fig. S15). Notably, the fibrillization process under 3 M urea occurred more rapidly in the presence of PEG8000 than in its absence, where fibrils appeared only after more than 48 hours (Fig. 2b and Supplementary Fig. S3). Similar condensate-associated fibrillar outgrowths have been reported for proteins such as FUS ^37^, hnRNPA1 ^13^, α-synuclein ^38^, and TDP-43 ^14^. No fibrillization was observed after 24 hours at 5 or 6 M urea (Supplementary Fig. S13). Thus, the molecular crowding shifts the regime favouring fibril formation upwards from 2 M urea to 3-4 M urea. Cy5-labelled PEG8000 did not enrich on the fibril, and the fluorescence remained uniformly inside the condensates, indicating that PEG8000 does not become a structural component of fibrils (Fig. 5e). However, it is interesting to note that after the addition of urea, the interior of the condensates dissolved, leaving fibrils in the dilute phase and in the shell of the condensates. This highlighted the different roles of the interior and the interface of the condensates in YPet-mR3 assembly, with the interface providing a favourable site for fibril development. ^17^

### The interface of YPet-mR3 condensates can be targeted to prevent fibrillization

It has been reported that the interface of biomolecular condensates can catalyse fibril formation. _39,40_ We therefore investigated whether modifying the interface of YPet-mR3 condensates could alter their fibrillization. YPet-mR3 was mixed with murine Z-DNA-binding protein RHIM domains fused to mCherry (mCh-mZBP1) (Fig. 6a). Murine Z-DNA binding protein senses z-form nucleic acids and recruits mRIPK3 via interactions between RHIMs, triggering RIPK3 self-phosphorylation and downstream necroptosis. Thus, addition of mCh-mZBP1 is expected to modulate the availability or conformation of the RHIMs and will likely impact the phase transitions of YPet-mR3. Unexpectedly, in the absence of a molecular crowder, mCh-mZBP1 coats the interface of YPet-mR3 condensates rather than partitioning uniformly throughout the condensates, even though these two proteins were added concurrently (Fig. 6b). We next tested whether the passivation of mCh-mZBP1 on YPet-mR3 condensates could prevent fibrillization under the crowding conditions at 3M and 4M urea. It was observed that mCh-mZBP1 formed a thin layer on the interface of YPet-mR3 condensates in the presence of NaCl. In the presence of CaCl_2_, mCh-mZBP1 penetrated further into YPet-mR3 condensates from the shell (Fig. 6c). The binding of mCh-mZBP1 was also accompanied by a reduction in the condensate size, especially for those that were incubated with CaCl_2_ (Supplementary Fig. S16a, b). Moreover, as previously described, YPet-mR3 formed fibrils at 3 and 4 M urea in the presence of a molecular crowder after 24-hour incubation (Fig. 5a). When mCh-mZBP1 was added, no fibrils were formed after 48 hours of incubation under the same conditions (Supplementary Fig. S16c). One possible explanation is that the interfacial recruitment of mCh-mZBP1 reduces productive homotypic interactions between YPet-mR3 molecules at the condensate interface and dilute-phase YPet-mR3, thereby preventing fibrillization. A related strategy has been demonstrated using a designed protein-based surfactant targeting the interface of hnRNPA1-B-LCD condensates, preventing their interface-induced fibrillization. ^13^ Consistent with an important role for the condensate interface, the complete removal of condensates can also prevent fibril formation over 48 hours (Fig. 6d).

**Figure 6.**
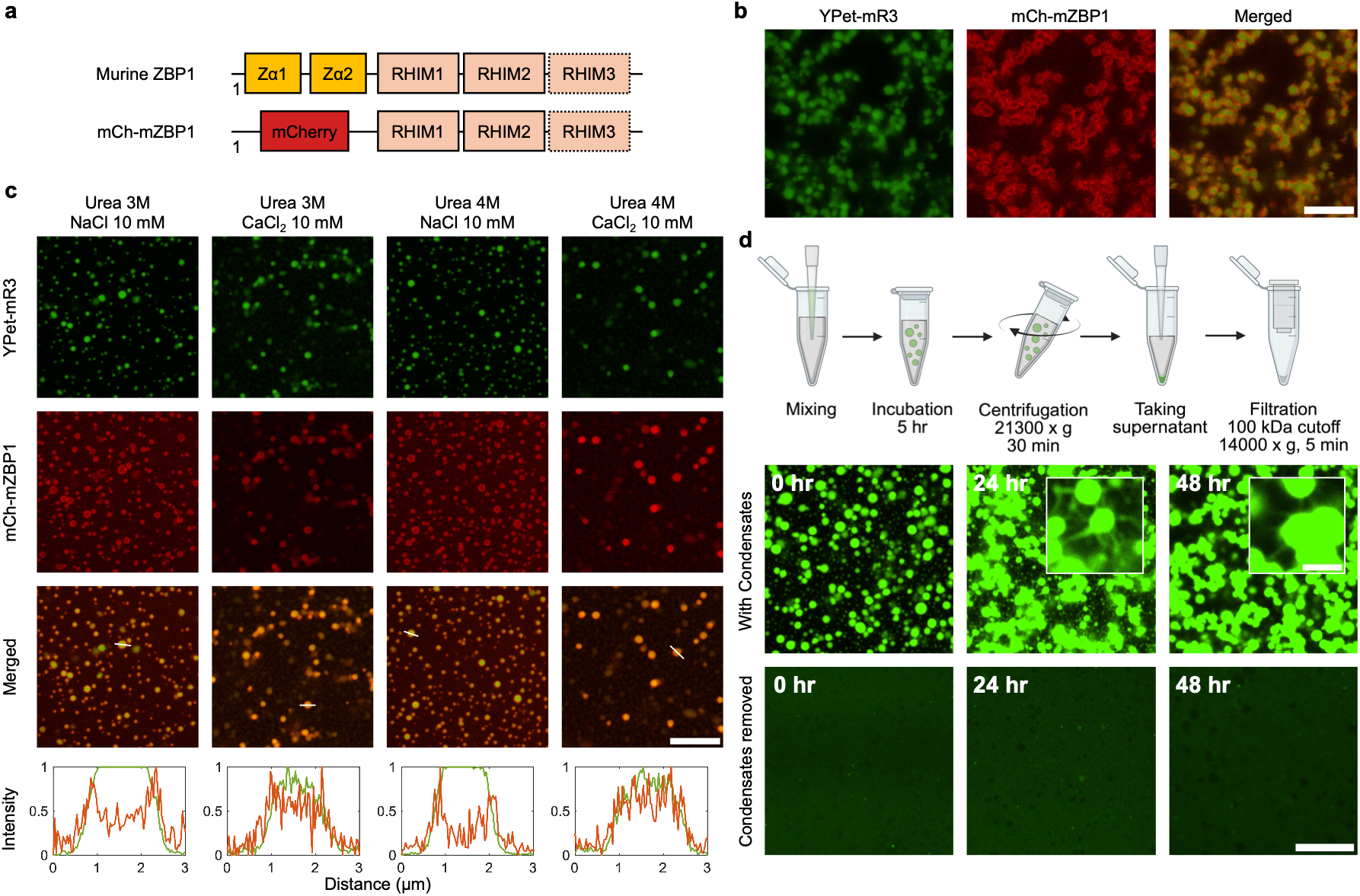
**a** Domain organisation of full-length murine ZBP1 and the mCh-mZBP1 construct used in these studies. **b** Confocal microscopic images 10 μM YPet-mR3 + 10 μM mCh-mZBP1 condensates prepared in Tris buffer with a final concentration of 0.8 M urea, 50 mM Tris-HCl, and 10 mM NaCl. Scale bar: 5 μm. **c** Confocal microscopic images 10 μM YPet-mR3 + 10 μM mCh-mZBP1 condensates incubated in Tris buffer in the presence of molecular crowder for 24 hours. The final solution includes 3 M/4 M urea, 50 mM Tris-HCl, 10 mM NaCl/CaCl_2_, and 10 wt% PEG8000. The YPet and mCherry intensity profiles (Green and Red) of the marked condensates are shown at the bottom. No fibrils were detected for these samples. Scale bar: 10 μm. **d** Centrifuge experiments for removal of 10 μM YPet-mR3 condensates in the presence of 3 M urea, 50 mM Tris-HCl, 10 mM NaCl, and 10 wt% PEG8000. No fibrils were detected within 48 hours after the condensates were removed. By contrast, fibrils developed from the interface of condensates. Scale bar: 10 μm. Inserts: 2 μm.

## Discussion and conclusions

Our findings revealed a LLPS-mediated pathway for YPet-mR3 phase transitions in parallel with the classical nucleation-elongation fibrillization pathway. The conformation of protein molecules determines the pathway of fibril formation and whether the process is promoted or suppressed. The formation of condensates does not simply affect the fibrillization in one form. Depending on the protein conformation and solution conditions, it can either kinetically retain YPet-mR3 in a reversible state or provide an interface that favours fibril nucleation and growth.

YPet-mR3 molecules form liquid condensates when they adopt a relatively folded conformation at low urea concentrations. Partially unfolded YPet-mR3 undergoes direct fibrillization without LLPS, whereas further unfolding at high urea concentrations keeps YPet-mR3 predominantly in the dispersed state. Intermediate urea concentrations can be optimal for amyloid formation, as they disrupt excessive attraction forces, partially unfold the protein, and expose amyloid-prone regions, thereby favouring amyloid formation. ^41,42^ In contrast, lower urea concentrations may permit or favour multivalent, transient interactions that disrupt the assembly into a well-ordered amyloid structure.

LLPS can be promoted by increasing the ionic strength. This shifts partially unfolded YPet-mR3 from direct fibrillization to the LLPS-mediated pathway and delays fibril formation. Ionic screening reduces repulsive forces between charged protein molecules and enhances homotypic interactions. ^42^ Salt containing divalent ions, such as CaCl_2,_ appears more effective than monovalent salts in promoting LLPS. We speculate that divalent ions may form salt bridges between negatively charged residues on protein molecules, as observed in silk fibroin condensates ^34^, although further work is required to obtain high-resolution binding information. Under several conditions, salt-induced LLPS remained reversible, in contrast with the corresponding fibrils formed without salt, indicating that condensates can serve as a kinetic trap that sequesters assembly-competent molecules and hinders their assembly into ordered amyloid structures.

The VQIG-to-AAAA mutation decouples the molecular requirements for LLPS from those for fibrillization. The mutant retained the ability to undergo LLPS and formed condensates over a broader range of urea concentrations, yet no fibrils were detected even after prolonged incubation. LLPS create a concentrated intermediate state, whereas the conversion from condensate to fibril requires a sequence-encoded amyloid switch that is not necessarily involved in condensate formation. The fibrillization is therefore influenced by how the aggregation-prone regions interact and undergo ordered structural conversion.

Molecular crowding further highlighted that the effect of LLPS on fibril formation is not uniform. PEG8000 elevates local concentration and drives YPet-mR3 molecules to form condensates under higher urea concentrations. Meanwhile, the molecular crowder reduces the volume available to protein molecules, lowers the configurational entropy of the unfolded states, and shifts the protein towards more folded states. ^32^ This explains why the PEG8000 fibrillization condition was upshifted from 2 M to 3-4 M urea. PEG8000 sequestered aggregation-prone YPet-mR3 molecules at 2 M urea into condensates, decreasing the fibrillization propensity by altering their conformation and delaying productive fibril assembly. When protein molecules adopt aggregation-prone conformations in the presence of 3-4 M urea and PEG8000, the large interfacial area provided by condensates can facilitate their deposition from the dilute phase, thereby promoting nucleation and subsequent fibrillization. ^43^ Passivating the condensate interface through addition of interaction partners such as ZBP1-RHIMs or removing the condensate effectively suppresses interface-induced fibrillization, indicating that condensate interfaces can be targeted to modulate fibrillization. ^44^

Our results extend the emerging framework by showing that the RIPK3-RHIM-containing construct can itself undergo LLPS and that this process can modulate amyloid fibril formation. However, the fluorescent tag, the isolated RHIM-containing construct, the denaturant, and the synthetic crowding conditions differ from those of the intracellular environment. The biological relevance of these assemblies will need to be further tested using full-length RIPK3 and cellular models. Nevertheless, this study serves as another example that condensates do not universally accelerate or delay fibrillization; instead, their effects depend on protein conformation, sequence-specific interactions, and physicochemical environment.

LLPS has been identified as a novel regulatory mechanism in innate immunity. ^45^ For future work, it would be interesting to determine how the RIPK3 kinase domain, post-translational modifications and interactions with RIPK1, TRIF, ZBP1 and other cellular components reshape its phase behavior. ^46,47^ It will also be important to establish whether condensates, fibrillar and non-fibrillar arrested assemblies differ in their ability to promote RIPK3 autophosphorylation, phosphorylate MLKL and trigger necroptotic cell death. Such studies could reveal whether phase separation acts as an upstream regulatory step in necroptotic signalling, a competing off-pathway state or a context-dependent intermediate that controls the kinetics and magnitude of RIPK3 activation.

## Methods

### Recombinant expression and purification of YPet-mR3, YPet-mR3-mut, and mCh-mZBP1

The His-tagged YPet or mCherry fusion proteins were produced recombinantly using a vector produced in-house. ^7^ Proteins were expressed in the BL21 (DE3) or Rosetta 2 strain of *E. coli* grown in ampicillin-containing LB media at 37 °C until an OD_600 nm_ of 0.6-0.8 was achieved. Expression was then induced with the addition of 0.5 mM IPTG and continued incubation at 37 °C for 3 hours.

Cell pellets were lysed via resuspension and sonication in lysis buffer (20 mM NaH_2_PO_4_, 150 mM NaCl, 2 mM EDTA, pH 8.0). The insoluble pellets obtained via centrifugation were resuspended in denaturing buffer (8 M urea, 20 mM Tris, pH 8.0) overnight at 4 °C. The sample was centrifuged at 16,000 g for 45 min at 4 °C, filtered and then purified on a 5-mL HisTrap column (GE Healthcare) under denaturing conditions using AKTA Start FPLC. The elution of protein was achieved with a gradient of 30-300 mM imidazole, monitored by absorbance at 280 nm.

The eluted fractions were analyzed by SDS-PAGE. The fractions containing the fusion proteins were collected and concentrated using Amicon Ultra-15 Centrifugal Filter Units with MWCO 10 kDa (Millipore).

### Phase transitions of YPet-mR3 and YPet-mR3-mut

YPet-mR3/YPet-mR3-mut stock solutions were stored in 8 M urea at 4 °C before use. The stock solution was diluted to 10 times the target protein concentration in 8 M urea to avoid a high concentration gradient. To trigger phase transitions, 0.5 μL of the stock solution was added to a 4.5 μL Tris buffer solution containing urea, Tris-HCl, and NaCl or CaCl_2,_ or PEG8000 10 wt% as applicable. For the fluorescent-tagged PEG8000 experiments, the final buffer consists of 1wt% of Cy5-PEG8000 (Biopharma PEG Scientific Inc.) and 9 wt% non-tagged PEG8000.

The mixture was stored in homemade PDMS-based well plates with glass slides as a base and sealed with Parafilm. The well plates were incubated at 25 °C in a mini-incubator (Cole-Parmer). All images were captured using a Nikon AX confocal microscope equipped with either a 60x water-immersed lens or a 100x oil-immersion lens.

### Fluorescence recovery after photobleaching

YPet-mR3 condensates with suitable sizes were prepared by diluting 200 μM YPet-mR3 in 8 M urea to 20 μM. The final solution contains 0.8 M urea, 10 mM NaCl, and 50 mM Tris-HCl (pH = 7.4). The condensates were incubated for different time periods, followed by photobleaching of a circular region of less than half of the condensates for 1 second, using a 488 nm laser line set to 10% laser power. The recovery was recorded for 3 minutes after bleaching using a Nikon AX confocal microscope with a 60x water-immersion lens. For YPet-mR3-mut, a 155 μM stock solution in 8 M urea was used, with other settings unchanged. The error bar was generated based on three individual measurements from different samples. The data were retrieved from NIS Elements (5.42.04) software, processed and plotted using MATAB (R2024b).

### Dissolution assay

For the dissolution assay without molecular crowder, YPet-mR3 at 100 μM was diluted to 10 μM in Tris buffer solution to final urea concentrations of 0.8 M, 1.4 M, 2 M, and 3 M. The final solution either contained no salts or contained 10 mM NaCl or 10 mM CaCl_2_. All samples included a final Tris concentration of 50 mM (pH = 7.4). Separate samples were prepared and incubated at 25 °C for 1, 5, 24, 48 hours, and 7 days. These samples were imaged using a Nikon AX confocal microscope equipped with a 60x water-immersion lens (pre-dissolution). Subsequently, the samples were mixed with 8 M urea at a 1:1 volume ratio and incubated at 25 °C for up to 5 hours before taking post-dissolution images.

For the dissolution assay in the presence of a molecular crowder, YPet-mR3 at 100 μM was diluted to 10 μM in Tris buffer solution to final urea concentrations of 0.8 M, 2 M, 3 M, and 4 M. The final solution either did not contain salts or contained 10 mM NaCl or 10 mM CaCl_2_. All samples included a final Tris concentration of 50 mM (pH = 7.4) and 10 wt% PEG8000. Separate samples were prepared and incubated at 25 °C for 1, 5, 24, 48 hours, and 7 days. These samples were imaged using a Nikon AX confocal microscope equipped with a 60x water-immersion lens (pre-dissolution). Subsequently, the samples were mixed with 8 M urea at a 1:2 volume ratio and incubated at 25 °C for up to 24 hours before post-dissolution images were taken.

### TEM

YPet-mR3 condensates and fibrils were prepared in Tris buffer as described. The condensates were loaded on carbon film-coated copper grids (CF400, emgrid, Australia) 10 minutes after LLPS without dilution. The grids were washed with Milli-Q water to prevent severe urea crystallisation. The fibrils were loaded onto carbon film-coated copper grids for 2 minutes, then negatively stained with 2% uranyl acetate for 10 seconds and washed with Milli-Q water. TEM images were taken using FEI Tecnai T12 TEM microscope with a voltage of 120 kV.

### Intrinsic Fluorescent Intensity measurement

The intrinsic fluorescent intensity of YPet-mR3 was measured using Horiba Fluoromax-4 spectrofluorometer. YPet-mR3 1 μM was diluted to 0.1 μM in a 50 mM Tris buffer (pH = 7.4) with final urea concentrations of 0.8 M, 2 M, and 3 M, respectively, containing or not containing 10 wt% PEG8000. The solution was loaded in a 10 mm quartz glass cuvette (Hellma). This concentration was used to achieve optimal intensity and prevent the scattering triggered by phase transitions. The samples were excited at 280 nm with an excitation slit of 5 nm. The spectra were collected from 300 nm to 400 nm with an increment of 1 nm, with the emission slit set to 5 nm. Blank buffer solutions with corresponding conditions were measured and subtracted. Four separate sets of samples were prepared and measured. The data points were fitted with a second-order Gaussian model and plotted using Matlab (2024b).

### Circular Dichroism spectroscopy

CD spectra were collected using Jasco J-810 polarimeter. YPet-mR3 150 μM was diluted to 15 μM in a 20 mM phosphate buffer (pH = 7.4) with final urea concentrations of 0.8M, 2M, and 3M, respectively. Spectra were acquired at 25 °C with 10 averaged scans from 250 to 210 nm at a scan rate of 20 nm min^−1^ using a 0.1 cm cuvette. Data were blank subtracted with the corresponding buffer solution.

### FTIR

FTIR spectra were collected using a Bruker Tensor 27 spectrometer with a Bruker BioATR II Unit. YPet-mR3 samples (10 μM YPet-mR3, 0.8M / 2M urea, 50 mM Tris-HCl) were directly prepared in the silicon sample well with a final volume of 15 μL, followed by covering with parafilm. A background of buffer solution with the same urea and salt conditions was subtracted. Measurements were made by averaging 128 scans at a resolution of 4 cm^−1^. The data were retrieved from OPUS (8.7.31) and plotted using MATLAB (2024b).

### SAXS

Small-angle X-ray scattering (SAXS) was performed in a co-flow setup on the SAXS/WAXS beamline at the Australian Synchrotron (Clayton, VIC). For each condition, at least 10 short exposures were averaged and radiation damage was excluded by frame-to-frame comparison; matched solvent blanks were subtracted to obtain the net scattering profiles. The sample-to-detector distance was 2.5 m (nominal *q* range 0.005–0.5 Å⁻¹); after removal of beamstop-affected and noise-dominated points the usable range was *q* ≈ 0.008–0.33 Å⁻¹ (condition-dependent; Supplementary Table S2). Because YPet-mR3 forms large, polydisperse assemblies, no classical Guinier region is present and all radii are reported as apparent values for relative comparison only. For the elongated and clustered assemblies we applied a cross-sectional Guinier analysis: ln[*q*·*I*(*q*)] was plotted against *q*² over a single fixed window (*q* = 0.012–0.035 Å⁻¹) common to all samples, and the apparent cross-sectional radius of gyration *R_c_* was obtained from the slope *m* as *R_c_* = √(−2*m*). *R_c_* is less sensitive than an overall *R_g_* to length polydispersity, finite rod length and weak low-*q* aggregation. Dimensionless Kratky plots were used only qualitatively (trends in peak height/position and low-*qR_g_* upturn) rather than as absolute indicators of folded versus disordered states. Real-space pair-distance distribution functions p(*r*) were obtained by indirect Fourier transform with GNOM/datgnom (ATSAS), ^48^ using the apparent *R_g_* of each curve as input so that *D*_max_ was set by a uniform rule across all samples; for the largest assemblies (0.8 M, and 2 M at the 10^th^ hour) *q*_min_·*D*_max_ exceeds π, so the corresponding *D*_max_ and p(*r*) tails are used only for relative comparison. Ab initio shape reconstructions were performed both with DAMMIF (ATSAS; P1 symmetry, independent runs averaged with DAMAVER) and, as an independent cross-check, with DENSS (v1.8.8; multiple independent maps aligned and averaged); the map-to-map Fourier shell correlation (FSC) resolution was used as a measure of reconstruction reproducibility (Supplementary Table S1). Envelopes were interpreted qualitatively, and the 0.8 M reconstructions (FSC ≈ 170–270 Å, comparable to the particle *R_g_*) are illustrative only.

### Determine the critical separation concentration via centrifugation

A standard curve based on fluorescent intensity was prepared by diluting stock YPet-mR3 to 0.1, 0.25, 0.5, 1, 2.5, and 5 μM in 8 M urea with a final volume of 20 μL. The fluorescent intensity was measured using a plate reader (CLARIOstar, BMG Labtech) with excitation at 508 nm and emission at 530 nm.

The critical separation concentration was determined using the methods described previously ^30^, with modifications tailored to the system. YPet-mR3 100 μM were diluted to 10 μM with a final urea concentration of 0.8-3M, Tris 50 mM, NaCl 10 mM, with or without PEG8000 10wt%. The samples were incubated for 5 hours to reach equilibrium. The samples were centrifuged at 25 °C, 21,300 g for 30 minutes using a benchtop centrifuge (Eppendorf). The supernatant was collected and filtered using 100 kDa MWCO centrifugal filters (Amicon) at 14,000 g for 5 minutes to remove residual condensates. 2 μL of the filtered solution was diluted 10-fold in 8 M urea, and the fluorescent intensity was measured using the microplate reader with the same settings as those used for the calibration curve. The data were retrieved from MARS and plotted using MATLAB (2024b).

### Co-assembly of mCh-mZBP1 and YPet-mR3

YPet-mR3 and mCh-mZBP1 stock solutions of 200 μM in 8 M urea were mixed in 1:1 ratio by volume. 0.5 μL of the mixture was added to 4.5 μL Tris buffer solution to a final solution containing 10 μM YPet-mR3 and 10 μM mCh-mZBP1 with the urea, salt and crowder conditions described. The samples were loaded in PDMS-based wells and sealed with parafilm. Confocal microscopic images were taken using a Nikon AX confocal microscope with 60x water-immersion lens. YPet channel was excited by a 488 nm laser, with an emission wavelength of 500-550 nm. mCherry channel was excited by a 561 nm laser with an emission wavelength of 606-721 nm. The size distribution of YPet-mR3 + mCh-mZBP1 and YPet-mR3 under the same buffer conditions was measured after 24 hours of incubation by image analysis using FIJI. To check the YPet-mR3 fibril formation, YPet-mR3 were overexposed by increasing the laser power.

### Condensate removal by centrifugation and filtration

YPet-mR3 100 μM was diluted to 10 μM in Tris buffer with a final urea concentration of 0.8 M, Tris 50 mM, and NaCl 10 mM. The sample was incubated for 5 hours to reach equilibrium, followed by centrifugation at 21,300 g for 30 minutes (Eppendorf). The supernatant was collected and filtered using 100 kDa MWCO centrifugal filters (Amicon) at 14,000 g for 5 minutes to remove residual condensates. The filtrate was loaded into PDMS-based wells and sealed with parafilm, followed by confocal microscopic imaging.

## Notes

### Competing Interest Statement

The authors have declared no competing interest.

### Funding

- Australian Research Council: DE230100837

