## Supporting Information for "A conformational switch governs the dual role of liquid–liquid phase separation in amyloid fibrillization"

### Urea and salt mediate YPet-mR3 phase transitions

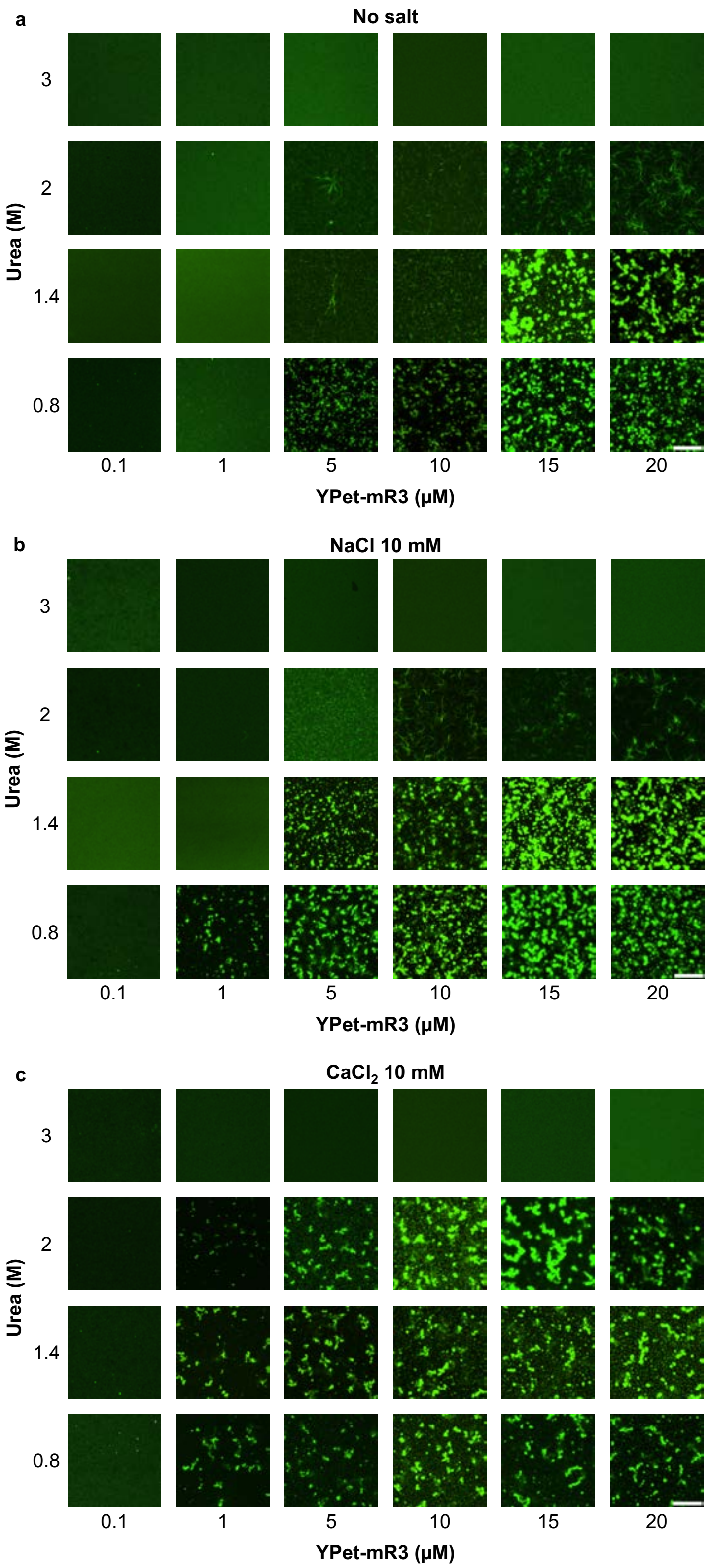


**Figure S1.** Phase diagram of YPet-mR3 (**a**) without salt, (**b**) with NaCl 10 mM, and (**c**) with $CaCl_2$ 10 mM. All final solutions include Tris 50 mM. All images were taken after 24 hours of incubation. Scale bar: 10 µm.

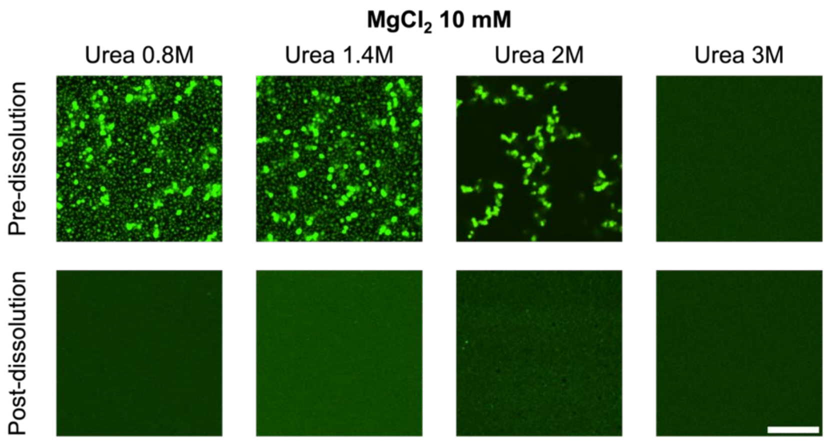


**Figure S2**. YPet-mR3 (10 µM) undergoes LLPS under 0.8 M – 2 M urea in the presence of $MgCl_2$, similar to $CaCl_2$. The formed condensates can be dissolved with 8 M urea after 24 hours of incubation. Scale bar: 10 µm.

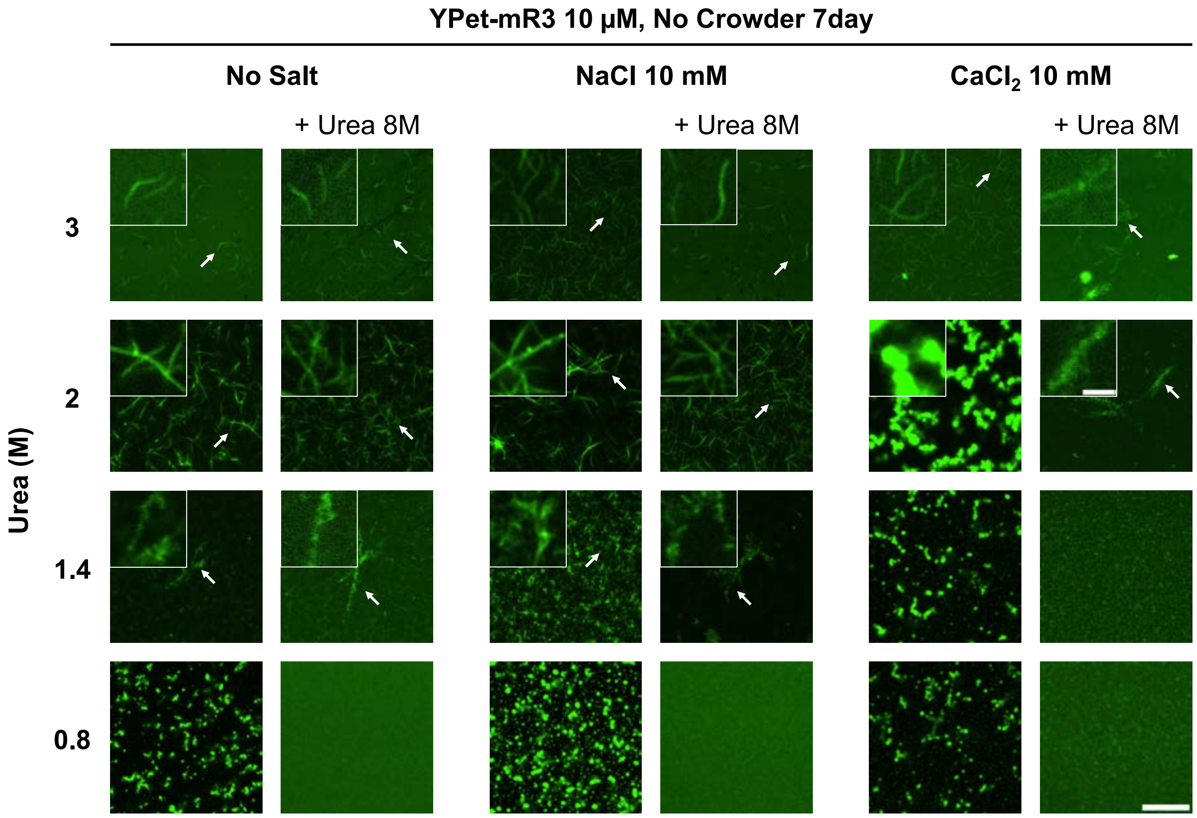


**Figure S3.** YPet-mR3 (10 µM) under different urea and salt conditions were incubated for 7 days, followed by adding 8 M urea for dissolution. Scale bar: 10 µm. Inserts: 2 µm.

### Conformational changes of YPet-mR3 at different urea concentrations

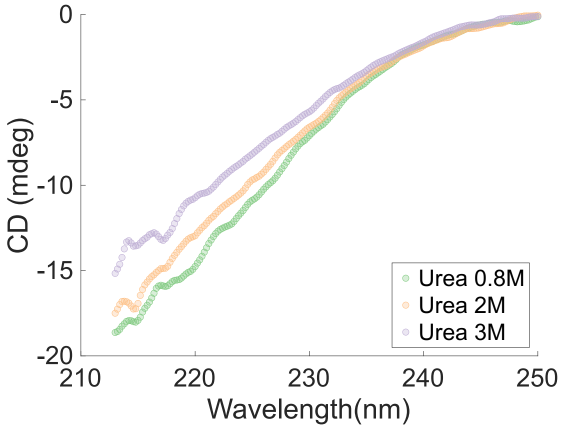


**Figure S4.** CD spectra of YPet-mR3 in the presence of 0.8 M, 2 M, and 3 M urea at $0^{th}$ hour.

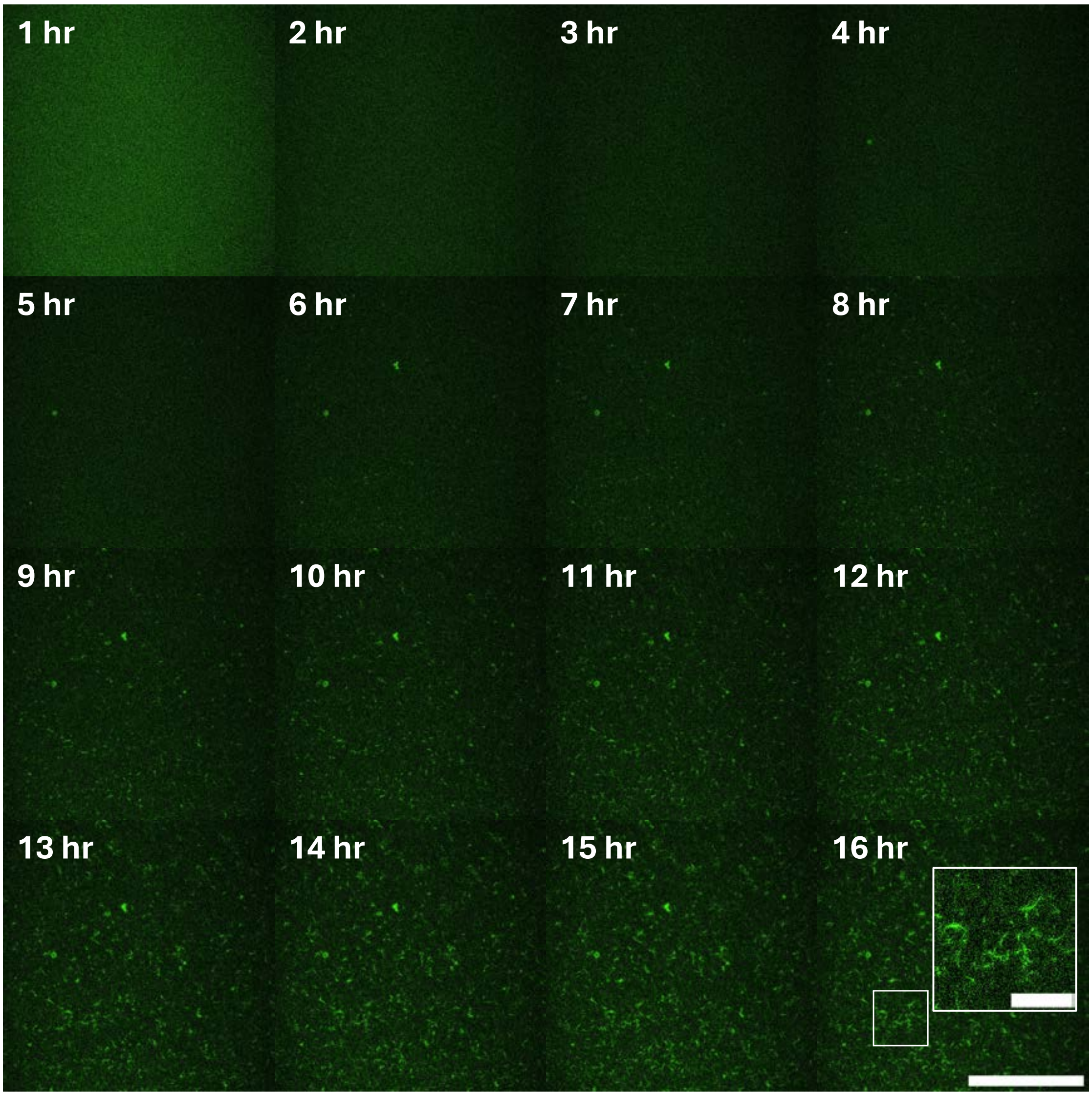


**Figure S5.** The fibrillation of YPet-mR3 10 μM under 2 M urea and Tris 50 mM. No LLPS was observed during the process. Scale bar: 50 μm, insert: 10 μm.

### Small-angle X-ray scattering (SAXS)

This section documents the SAXS analysis underlying Figure 3d-f of the main text. For all samples, YPet-mR3 self-associates into large, polydisperse assemblies, so the scattering contains no classical Guinier region; every size reported here is an apparent, intensity-weighted average, valid for comparison between conditions and time points but not as the radius of a single monodisperse particle. Four complementary analyses are presented.

First, the cross-sectional (rod) Guinier analysis (Figure S6) measures the effective thickness of the assemblies. Plotting $\ln[q\cdot I(q)]$ against $q^2$ gives a slope m from which the apparent cross-sectional radius $R_c = \sqrt{(-2m)}$ is obtained. The fit uses a single window (q = 0.012–0.035 $Å^{-1}$) identical for every sample, so differences in $R_c$ reflect the samples rather than the fitting choice. $R_c$ is preferred over an overall $R_g$ because it is far less sensitive to length polydispersity, finite rod length and weak interparticle scattering at the lowest q. Dimensionless Kratky plots (Figure S7), used only qualitatively, show the change in the low-$qR_g$ region under 2 M and 3 M urea.

Second, the pair-distance distribution p(r) (Figure S8) converts the same data into real space. A practical point determined the quality of every p(r) here: because p(r) is a low-resolution quantity, the indirect transform was restricted to $q \leq 0.10$ $Å^{-1}$ ($\approx 8/R_g$). Using the full measured range let the noisy high-q tail dominate and produced strongly oscillating distributions; with the restricted range the oscillation criterion improved from 3.6–6.3 to 1.13–1.26 for the 0.8 M series and the GNOM total estimate rose from 0.54–0.62 to 0.75–0.88. Dmax was then chosen by scanning it and taking the solution with the lowest oscillation among those that are strictly positive and approach zero smoothly.

The quality of each p(r) can be judged from the criteria GNOM reports (Table S1). All twelve distributions are strictly positive (POSITV = 1.00) and the oscillation criterion is 1.13–2.03 against an ideal 1.1. Two limitations remain. For the 0.8 M clusters qmin·Dmax = 5.1–6.2 exceeds π, so Dmax and the large-r part of p(r) are not resolved by the measurement and are used only as relative indicators. At 3 M urea the scattering is weakest and the transform still rings slightly (OSCILL 1.6–2.0), so features narrower than about 50 Å are not interpreted.

Third, ab initio shape reconstruction was performed independently with two algorithms: DAMMIF, which builds a dummy-atom bead model, and DENSS, which reconstructs an electron-density map without assuming uniform density (Figures S9 and S10). Agreement between two methods with different assumptions is stronger evidence than either alone. Reproducibility is quantified by the number of DAMAVER clusters among six independent runs and by the map-to-map Fourier shell correlation (FSC) resolution of ten DENSS maps.

Fourth, Figure S11 shows how well each model reproduces the measured curve. With the corrected input, all twelve datasets are fitted well by both methods (DAMMIF $\chi^2$ 0.83–1.29; DENSS $\chi^2$ 0.74–1.77), so poor fitting is no longer a limitation for any condition. What does limit interpretation is reproducibility. For the 0.8 M clusters the DENSS FSC resolution (211–261 Å) is comparable to the particle $R_g$ (229–272 Å), i.e. only the overall size — not the internal shape — is determined by the data, and these envelopes are shown for illustration only. The 2 M and 3 M reconstructions reach 48–110 Å, well below their $R_g$, and describe genuine low-resolution shapes. Widening the modelling q-range was tested and rejected: it lowered $\chi^2$ but worsened the FSC resolution in seven of eight samples, the signature of overfitting.

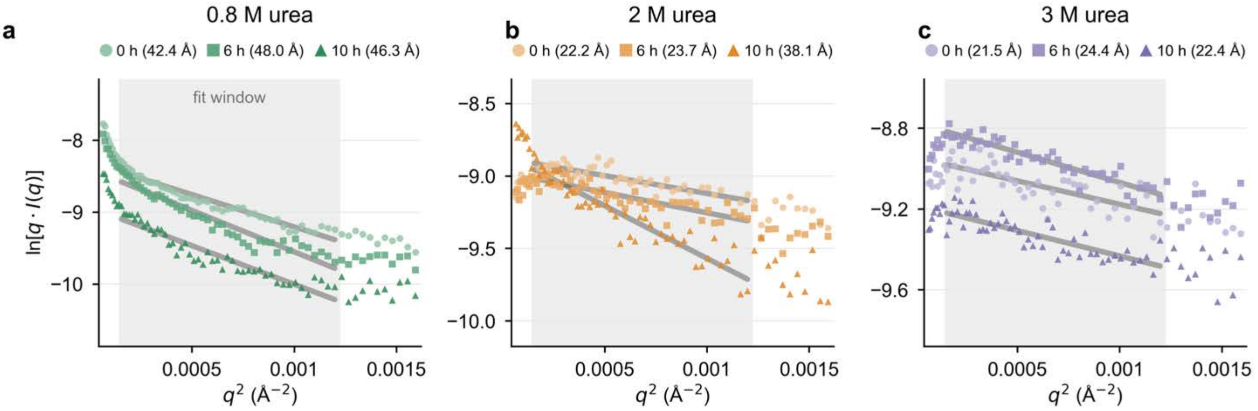


**Figure S6.** Cross-sectional (rod) Guinier plots, $\ln[q \cdot I(q)]$ versus $q^2$, for YPet-mR3 at (a) 0.8 M, (b) 2 M and (c) 3 M urea at 0, 6, and 10 hours. Shading marks the fixed fit window ($q = 0.012$–$0.035$ Å$^{-1}$) used for every sample; grey lines are the corresponding linear fits, from whose slope m the apparent cross-sectional radius $R_c = \sqrt{(-2m)}$ is obtained. Fitted $R_c$ values are given in the panel legends. The y-scale differs between panels, so quantitative comparison should use the $R_c$ values rather than the apparent steepness of the lines.

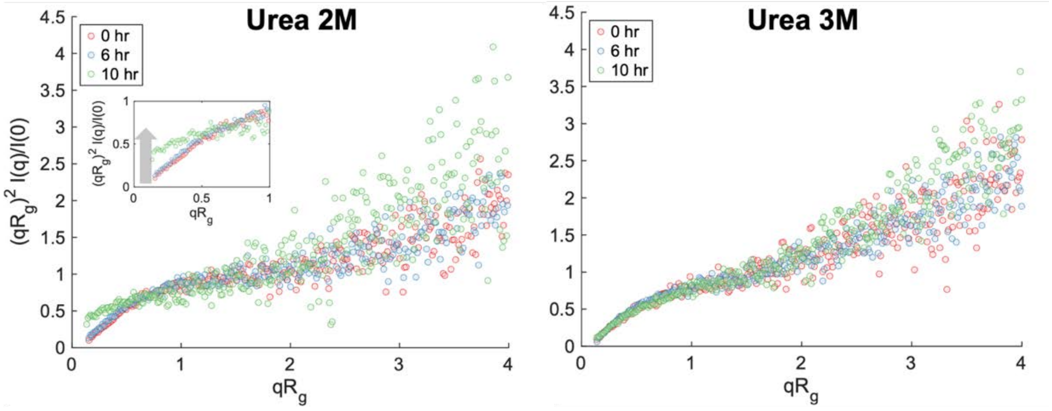


**Figure S7.** Dimensionless Kratky plots of YPet-mR3 in the presence of 2 M and 3 M urea over time.

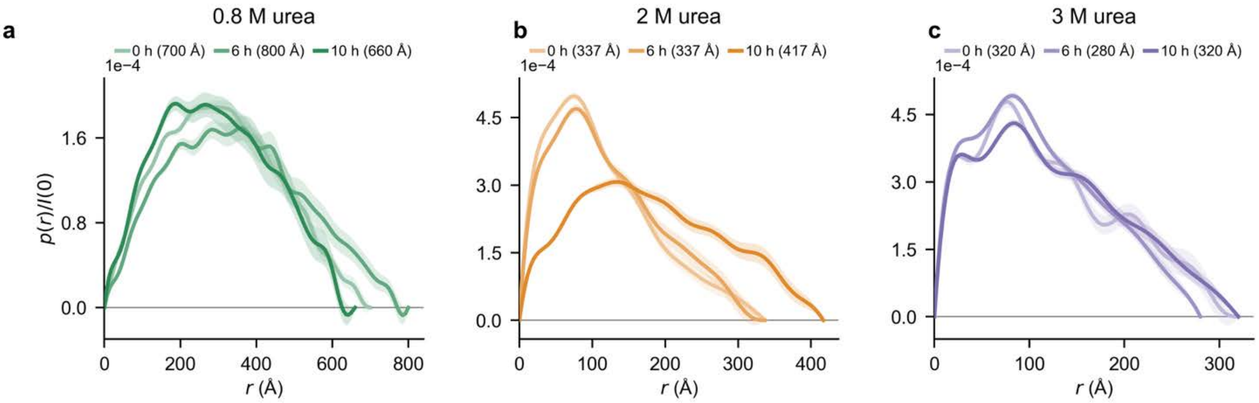


**Figure S8.** Pair-distance distribution functions p(r), normalised to I(0), for (a) 0.8 M, (b) 2 M and (c) 3 M urea at 0, 6, and 10 hours, with the selected Dmax given in each legend and the GNOM uncertainty shown as a shaded band. All transforms used $q \leq 0.10$ Å$^{-1}$ and a Dmax selected by scanning (Table S1). The 0.8 M distributions are broad and essentially unchanged with time; at 2 M the distribution broadens markedly at 10 h (Dmax 337 → 417 Å) and then contracts again by 19 hours; at 3 M it remains narrow throughout.

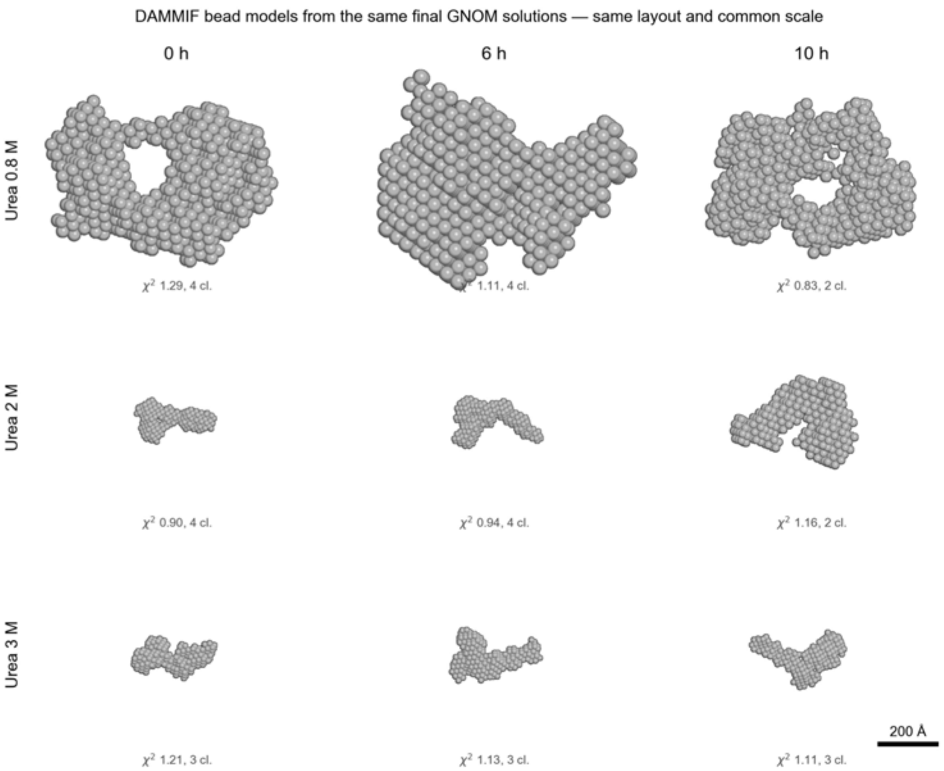


**Figure S9.** DAMMIF dummy-atom bead models (six independent runs averaged with DAMAVER) for the same conditions and time points, with the same layout and common scale as Figure S8; $\chi^2$ and the number of DAMAVER clusters are annotated per panel. This independent reconstruction method reproduces the trend seen with DENSS.

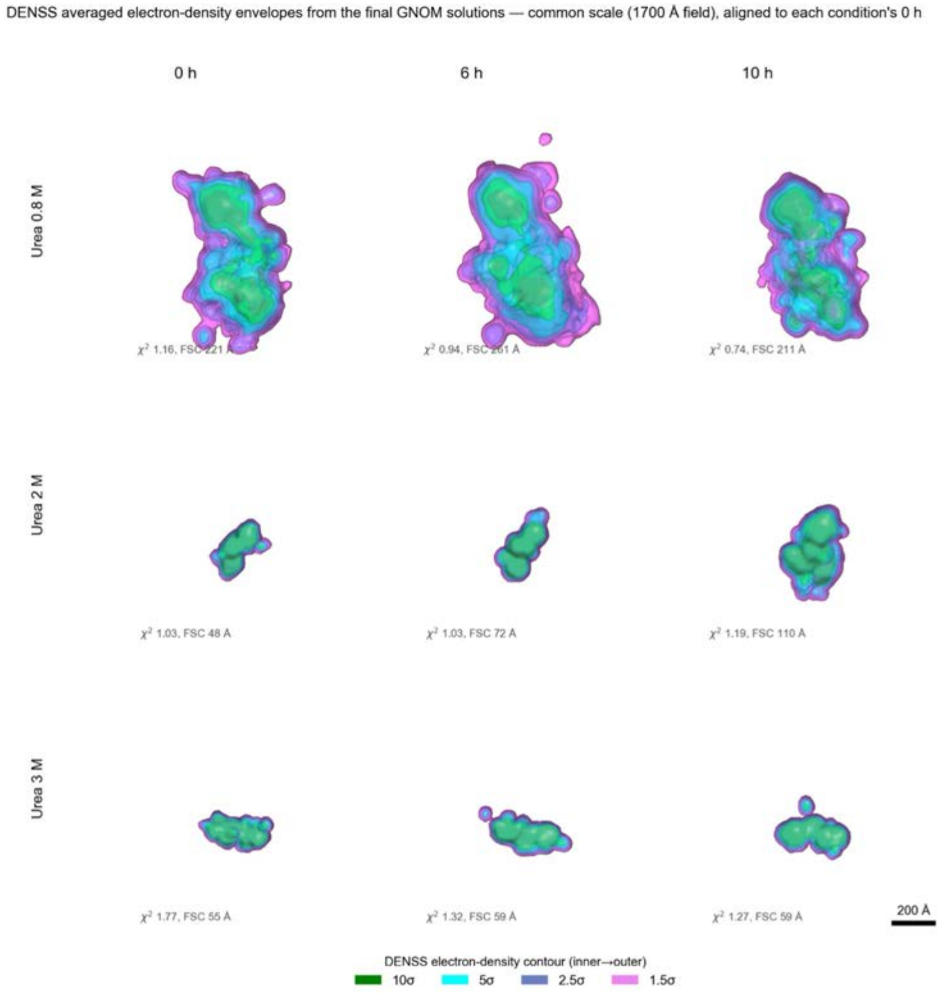


**Figure S10.** DENSS averaged electron-density envelopes at 0, 6, and 10 hours, all panels at the same scale (1700 Å field) and aligned to each condition's 0 h reconstruction. Four nested contour levels (1.5σ, 2.5σ, 5σ, 10σ) are shown; $\chi^2$ and FSC resolution are annotated per panel. The 0.8 M assemblies are the largest at every time point but their shape is not uniquely determined (FSC 211–261 Å, comparable to the particle Rg) and is shown for illustration only.

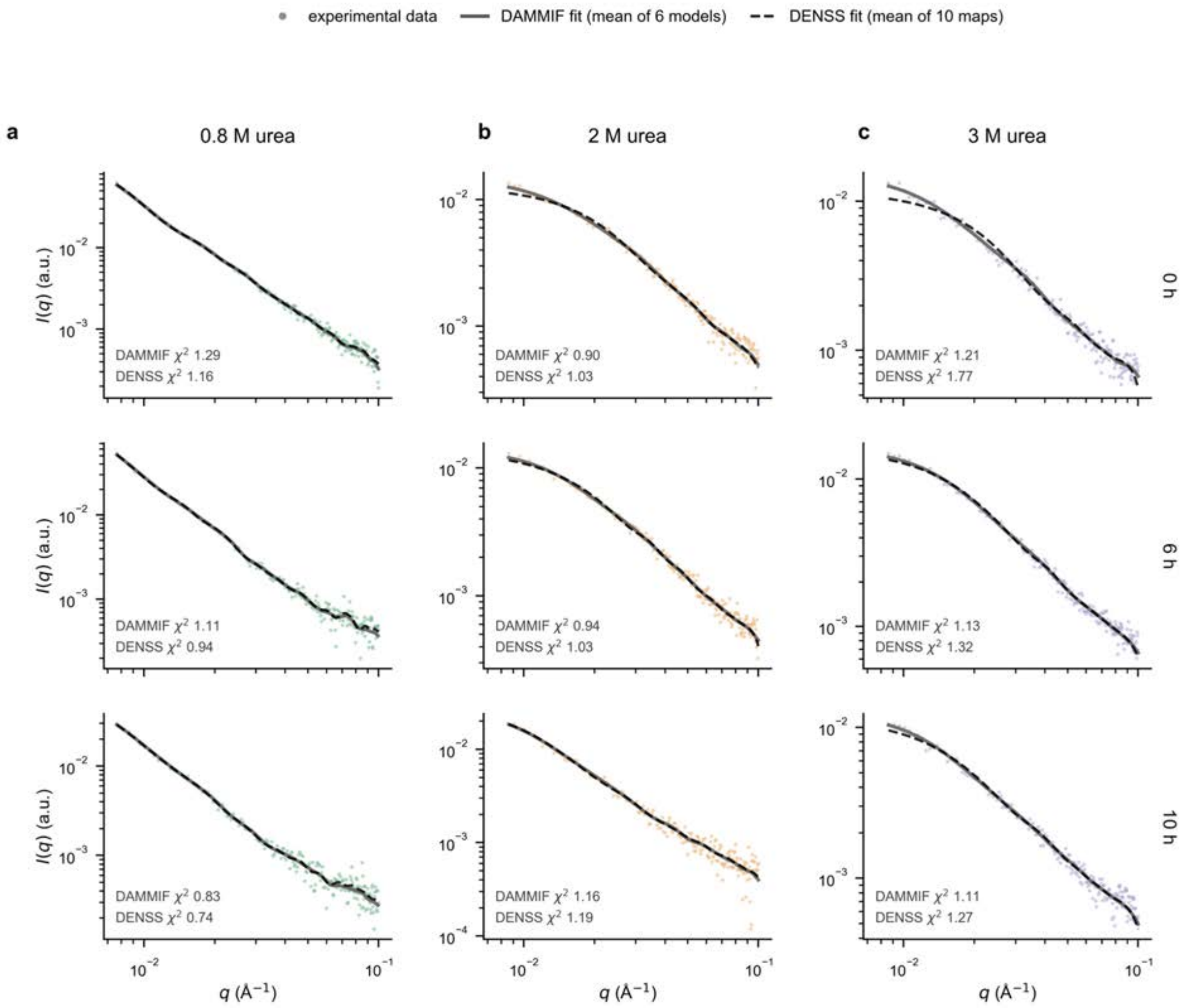


**Figure S11.** Fits of the two ab initio models to the experimental scattering curves for (a) 0.8 M, (b) 2 M and (c) 3 M urea at 0, 6, and 10 hours(log–log). Points are the experimental data. The grey solid line is the DAMMIF fit (mean of six models) and the black dashed line the DENSS fit (mean of ten maps); $\chi^2$ values are annotated in each panel. Both methods describe all twelve datasets well.

### Table S1. SAXS fitting and modelling parameters

All values were obtained from the analyses in Figures S6–S10. A single cross-sectional Guinier window and a uniform p(r) protocol ($q \le 0.10$ Å$^{-1}$, Dmax selected by scanning) were used for every sample.

| Urea | Time | Rc (Å) | max q·Rc | $R^2$ | Dmax (Å) | Rg (Å) | qmin·Dmax | OSCILL | GNOM est. | DAMMIF χ² | clusters | DENSS χ² | FSC (Å) |
|---|---|---|---|---|---|---|---|---|---|---|---|---|---|
| 0.8 M | 0 h | 42 ± 1 | 1.47 | 0.95 | 700 | 241 ± 4 | 5.39 * | 1.13 | 0.80 | 1.29 | 4/6 | 1.16 | 221 |
| 0.8 M | 6 h | 48 ± 1 | 1.66 | 0.93 | 800 | 272 ± 7 | 6.16 * | 1.26 | 0.75 | 1.11 | 4/6 | 0.94 | 261 |
| 0.8 M | 10 h | 46 ± 1 | 1.60 | 0.91 | 660 | 229 ± 7 | 5.08 * | 1.22 | 0.88 | 0.83 | 2/6 | 0.74 | 211 |
| 2 M | 0 h | 22 ± 1 | 0.77 | 0.72 | 337 | 94 ± 3 | 2.90 | 1.84 | 0.50 | 0.90 | 4/6 | 1.03 | 48 |
| 2 M | 6 h | 24 ± 1 | 0.82 | 0.78 | 337 | 99 ± 4 | 2.90 | 1.71 | 0.76 | 0.94 | 4/6 | 1.03 | 72 |
| 2 M | 10 h | 38 ± 1 | 1.32 | 0.89 | 417 | 144 ± 3 | 3.59 * | 1.33 | 0.64 | 1.16 | 2/6 | 1.19 | 110 |
| 3 M | 0 h | 22 ± 1 | 0.74 | 0.55 | 320 | 99 ± 3 | 2.75 | 2.03 | 0.45 | 1.21 | 3/6 | 1.77 | 55 |
| 3 M | 6 h | 24 ± 1 | 0.85 | 0.87 | 280 | 90 ± 1 | 2.41 | 1.61 | 0.59 | 1.13 | 3/6 | 1.32 | 59 |
| 3 M | 10 h | 22 ± 1 | 0.78 | 0.70 | 320 | 102 ± 2 | 2.75 | 1.86 | 0.65 | 1.11 | 3/6 | 1.27 | 59 |

Cross-sectional Guinier: fixed window $q = 0.012–0.035$ Å$^{-1}$; $R_c = \sqrt{(-2m)}$ with the error propagated from the slope; strictly valid for $q \cdot R_c < 1.3$. p(r): GNOM with $q \le 0.10$ Å$^{-1}$; Dmax selected by scanning and taking the lowest OSCILL among strictly positive solutions that approach zero smoothly (ideal OSCILL 1.1); * marks qmin·Dmax > π, i.e. Dmax beyond the reliably resolved range. DAMMIF: fast mode, P1, six independent runs; 'clusters' is the number of DAMAVER clusters among them (one cluster = fully reproducible). DENSS v1.8.8: ten independent maps aligned and averaged; FSC is the map-to-map Fourier shell correlation resolution at 0.5.

### Table S2. SAXS sample, data-collection and analysis parameters

Reported following the recommendations for publishing biomolecular small-angle scattering experiments (Trewhella et al., Acta Cryst. D 2017).

| **Parameter** | **Value** |
|---|---|
| (a) Sample | |
| Protein / construct | YPet-mR3 (YPet fused to the murine RIPK3 RHIM); ~317 residues, ~35 kDa monomer |
| Concentration | 10 µM (~0.35 mg $mL^{-1}$) |
| Solvent | 50 mM Tris-HCl pH 7.4 with 0.8 M, 2 M or 3 M urea |
| Temperature | 25 °C |
| Sample state | 0.8 M: phase-separated; 2 M: fibrillating; 3 M: dispersed |
| (b) Data collection | |
| Beamline / instrument | SAXS/WAXS beamline, Australian Synchrotron (Clayton, VIC); co-flow cell |
| Sample–detector distance | 2.5 m |
| q range (nominal / measured) | 0.005–0.5 $Å^{-1}$ / 0.0053–0.48 $Å^{-1}$ |
| q range used for p(r) | ≤ 0.10 $Å^{-1}$ (≈ 8/Rg; p(r) is a low-resolution quantity) |
| Exposures | ≥10 short frames averaged per condition |
| Radiation damage | Excluded by frame-to-frame comparison |
| Background subtraction | Matched solvent blanks (zero-background reduction) |
| (c) Analysis and modelling | |
| Cross-sectional Guinier | ln[q·I(q)] vs $q^2$, fixed window q = 0.012–0.035 $Å^{-1}$; $R_c = \sqrt{(-2m)}$ |
| Kratky analysis | Dimensionless Kratky, used qualitatively |
| p(r) / indirect Fourier transform | GNOM (ATSAS 3.2.1); Dmax selected by scanning (see Table S1) |
| Ab initio bead modelling | DAMMIF (ATSAS 3.2.1), fast mode, P1, 6 runs, averaged with DAMAVER |
| Ab initio density modelling | DENSS v1.8.8, 10 independent maps, aligned and averaged |
| Reproducibility metrics | DAMAVER cluster count; DENSS map-to-map FSC (0.5) resolution |
| Rendering | PyMOL (open-source); density maps at 1.5σ, 2.5σ, 5σ and 10σ |
| (d) Data availability | |
| Deposition | Scattering data and models to be deposited in SASBDB |

### The role of core tetrad residues on YPet-mR3 phase transitions

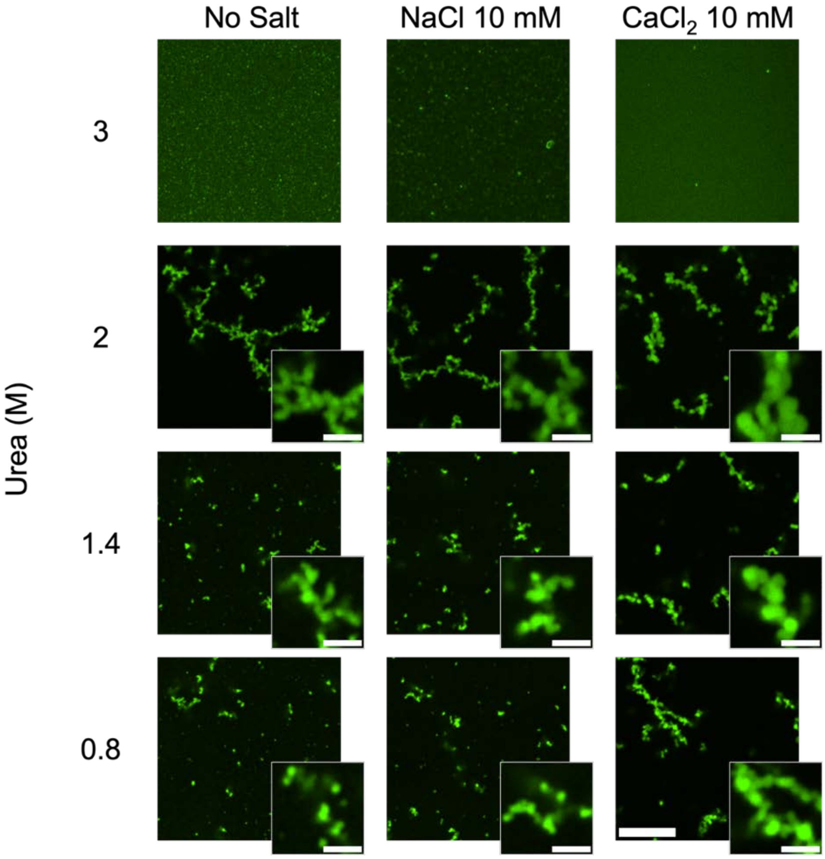


**Figure S12. a** Confocal microscopic images of 10 µM YPet-mR3-mut incubated under 0.8, 1.4, 2, 3 M urea in Tris-HCl 50 mM for 7 days. Scale bar: 10 µm, insert: 2 µm.

### Modulating the phase behavior of YPet-mR3 by using molecular crowder

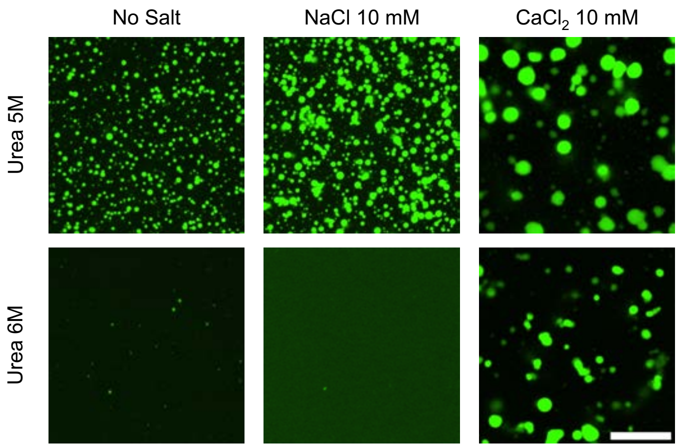


**Figure S13.** YPet-mR3 condensate incubated in the presence of 10 wt% PEG8000 and 5 M / 6 M urea with different salt conditions after 24 hours. Scale bar: 10 µm.

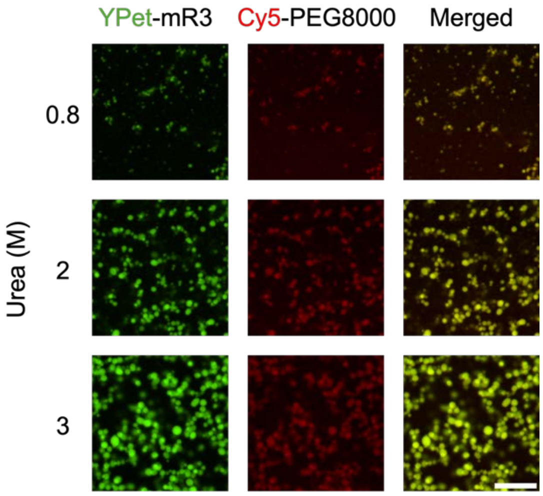


**Figure S14.** Confocal microscopic images of 10 µM YPet-mR3 in the presence of 10wt% PEG8000 (1wt% Cy5-PEG8000 + 9wt% PEG8000) and 50 mM Tris-HCl. Scale bar: 10 µm.

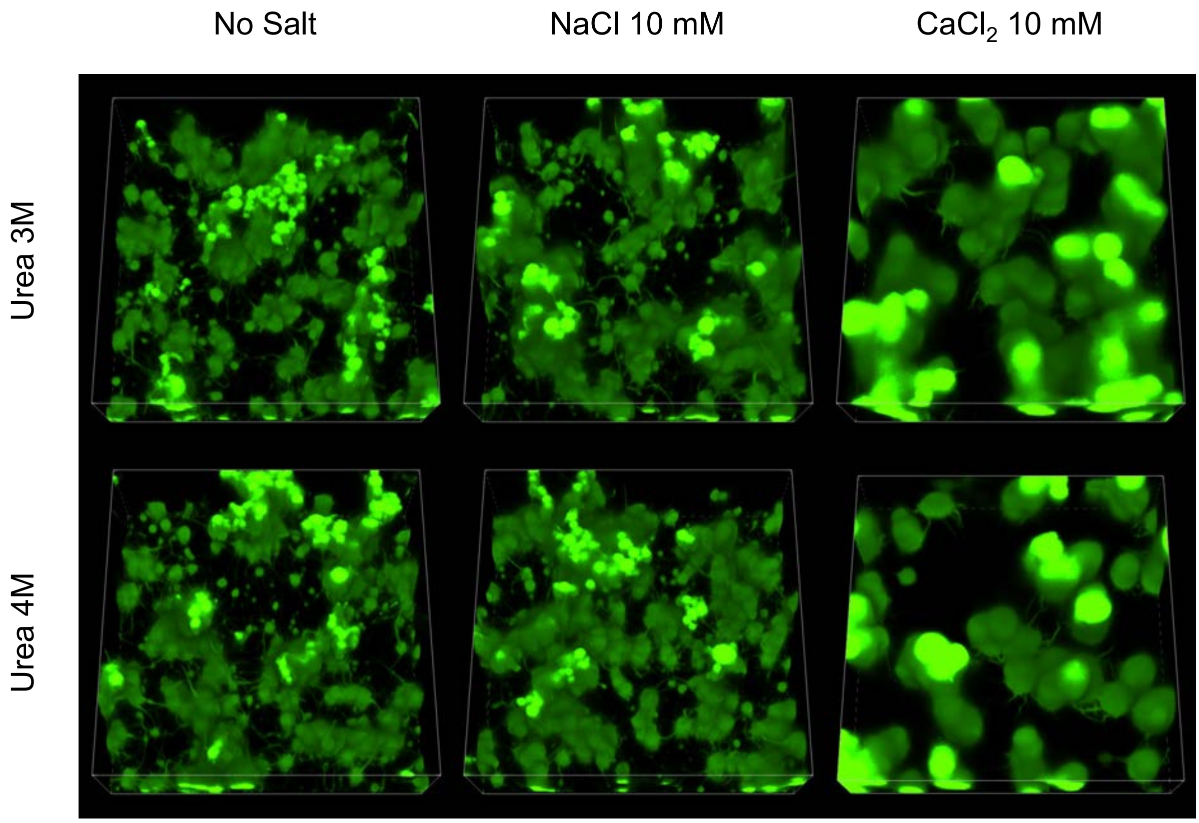


**Figure S15.** Z-scan of YPet-mR3 condensates incubated in the presence of 10 wt% PEG8000 under different urea and salt conditions after 48 hours.

### The interface of YPet-mR3 can be targeted to prevent fibrillization

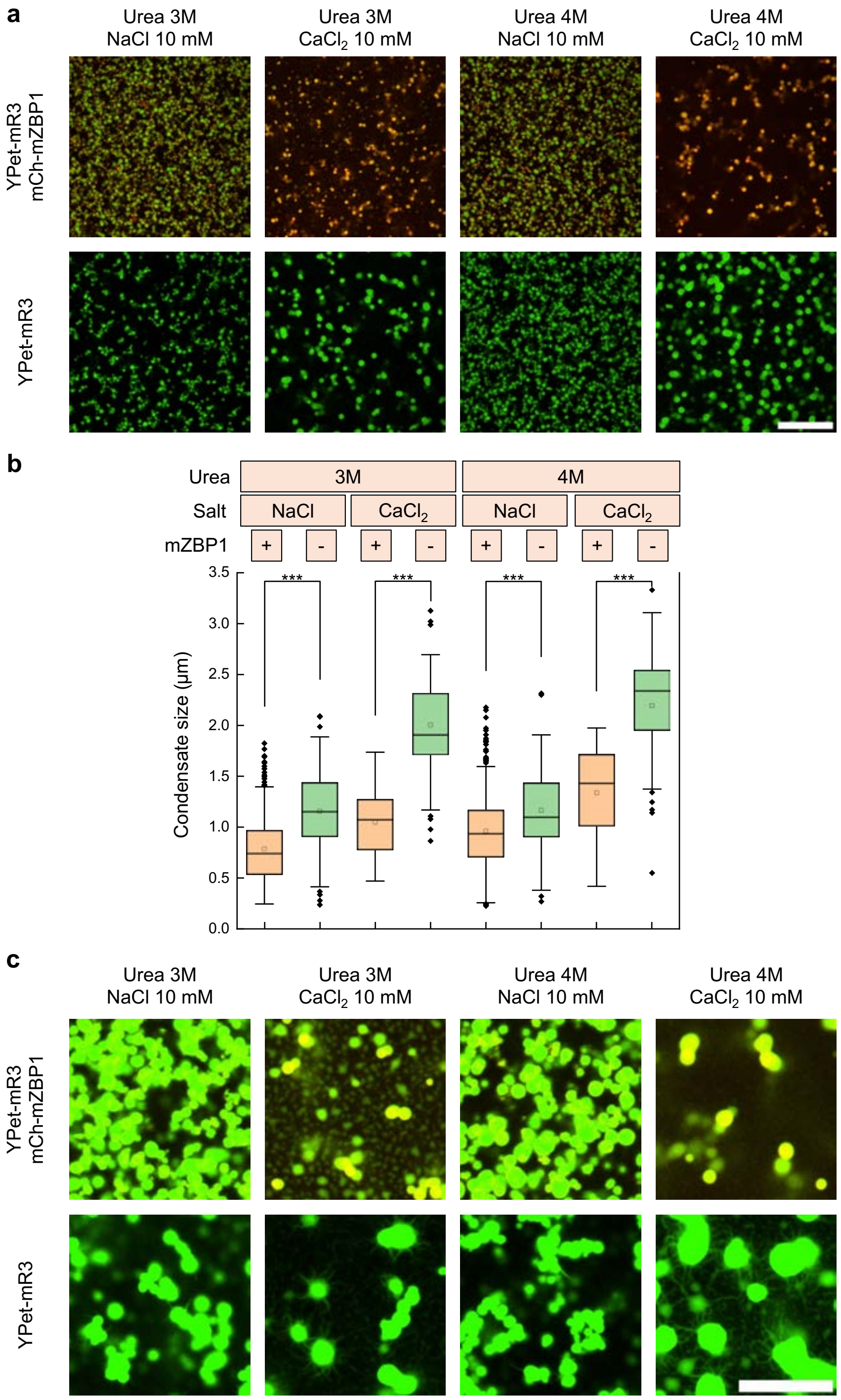


**Figure S16**. **a** Confocal microscopic images of 10 μM YPet-mR3 + 10 μM mCh-mZBP1 condensates and 10 μM YPet-mR3 condensates incubated for 24 hours under different conditions (3 M/4 M urea, 50 mM Tris-HCl, 10 mM NaCl/CaCl2, and 10 wt% PEG8000). Scale bar: 20 μm. **b** Size distributions of condensates. Paired t-tests were conducted. *** $p < 0.001$. **c** Confocal microscopic images of YPet-mR3 10 μM + mCh-mZBP1 10 μM and YPet-mR3 10 μM incubated in the presence of urea 3 M / 4 M, Tris 50 mM, NaCl / CaCl2 10 mM, PEG8000 10wt% for 48 hours. The YPet channel is overexposed to visualize the fibril formation. Scale bar: 10 μm.
